# Evaluation of nutraceuticals including vitamin B_12_ and xenobiotic degradation capacity of *Pleurotus* species

**DOI:** 10.1101/805382

**Authors:** Yashvant Patel, Sanjay Kumar Vishwakarma, Kumari Sunita

## Abstract

Diverse edaphic zone (namely: usar, wastelands, forest area, wetlands, flood prone area and fertile lands) were identified in the eastern part of Uttar Pradesh and collected species of *Pleurotus* for present study. A total of 39 purified isolates were evaluated for the presence of neutraceuticals like proteins, carbohydrates, phenolic and vitamin B_12_ along with xenobiotic dye degradation capacity for textile dyes (MG and BPB) and production of laccase enzyme simultaneously. Isolate no. 06, appeared most distant in dendrogram having two major clusters, which also showed highest MG degradation capacity, however, other isolates also showed excellent degradation of BPB; and the laccase activity was found in the range of 4.03 to 19.13 IU/ml of crude enzyme extract from mycelia. All the isolates used in the present study, were also mounted for their genetic diversity analysis through RAPD. Diversity study revealed gene frequency from 0.012 to 0.987 and the average gene diversity for all RAPD loci were 0.244. The Shannon Information Index was 0.397. The unbiased genetic similarity among all pairs of isolates was 0.36 to 0.93 with a mean of 0.64. Significant genetic diversity, nutraceuticals and laccase enzyme availability and dye degradation capacity within the studied genus *Pleurotus* was found, which makes necessary to carry out a selection process in each one for superior selection not only for human being but also many aquatic as well as other terrestrial flora and fauna. Present investigation suggest that due to wide range of variation within species, the adaptation of strains to different edaphic zones must be taken into account in selection processes.

## 1. INTRODUCTION

The genus *Pleurotus*, belongs to class Basidiomycetes (Castellano, 1989), is one of the most important edible mushrooms along with *Agaricus, Volvariella* and *Lentinula*, and possess many bio-potentialities right from nutritive mushroom production to bio-degradative non-specific extra cellular enzyme production. This genus has pool of non-specific enzymes (e.g. laccases, polyphenol oxidase, xylanase) that help in the efficient colonization and decomposition of a wide variety of lignocellulosic materials. These mushrooms play a substantial role in degradation of xenobiotic compounds mainly textile dyes due to many non-specific extra cellular enzymes (Rajak et al., 2011; Royse, 2002; Hofrichter, 2002). Along with these lignocellulosic degradative properties, it has been shown to possess many important bio-potentialities together with various environmental and biotechnological applications (Rajarathnam and Bano, 1989; Rajarathnam et al., 1992, 1998; Vishwakarma et al., 2012) including degradation of xenobiotics (Young et al., 2015; Loi et al., 2016). It also plays critical role in human health, agriculture and food industry and as model organisms for basic scientific studies (Valverde et al., 2015; Patel et al., 2012, 2017).

Traditionally, edible species of the genus *Pleurotus* were considered as medicinally important mushrooms (Bano and Rajarathnam, 1988; Khan and Tania, 2012). Now, it has been proved by many studies that this mushroom possesses many medicinally important properties like antibacterial, antiviral, immunomodulatory, hypocholesterolemic, anticholesterolic, antimutagenic, hyperglycemic etc. as reviewed by many researchers (King, 1993; Patel et al., 2012; Valverde et al., 2015; Gregori et al., 2007). It also possesses many important nutritional supplements like vitamins, fibres, proteins and essential amino acids and low cholesterols (Mattila et al., 2001). Because of all these properties *Pleurotus* spp is considered as “nutraceuticals” (Chang and Buswell, 1996).

Currently, commercial mushroom production is based on the limited number of strains available, which are at high risk of environmental changes; and hence genetic diversity is critical for adaptation to environmental changes and for the long-term survival of the species. Knowledge of genetic diversity within and among populations has practical importance for conservation and management policies (Hamrick and Godt, 1996; Fritsch and Rieseberg, 1996). The preservation of genetic diversity within the species is a major target of conservation, because loss of genetic variation is thought to reduce the ability of populations to adapt to environmental change for survival (Fisher et al., 2012; Parmesan, 2006; Hogbin and Peakall, 1999). Therefore, population genetic studies of *Pleurotus* spp are essential for providing necessary information for conservation of this very important genus.

PCR-based DNA fingerprinting techniques such as Random Amplified Polymorphic DNA analysis (RAPD), Inter Simple Sequence Repeat (ISSR) and Amplified Fragment Length Polymorphism (AFLP) represent a very informative and cost-effective approach for assessing genetic diversity for a wide range of organisms. All these markers do not require any prior knowledge of the genome of the species (Williams et al., 1990; Zietkiewicz et al., 1994; Bornet and Branchard, 2001). RAPD has been the most employed technique because it is simple and fast. Despite questions about its reproducibility, its utility in diversity analysis, mapping and genotype identification has been exploited in many plant species (Jone et al., 1997).

The species diversity of genus *Pleurotus* at DNA level is completely lacking in the studied area. This study by the strain identification and genetic analysis of the *Pleurotus* spp could help to uncover novel, economically important genetic variations for breeding purposes and removal of pollutants from water bodies and agro-industrial waste materials. Studying the genetic diversity using RAPD markers provide an opportunity to scan the entire genome for direct comparison of genetic materials that is almost independent of environmental influences (Zhao and Pan, 2004; Bautista et al., 2003; Harvey and Botha, 1996). This is the first study on inter-population genetic diversity of *Pleurotus* in this diverse geographical region. In the present work, there were three main objectives: (i) collection of the species of *Pleurotus* from different edaphic zones and evaluate neutraceuticals variability i.e. proteins, carbohydrates, vitamins, phenolic content and laccase activity, (ii) evaluation of textile dye degradation potential of all the collected isolates of *Pleurotus* spp, (iii) evaluation of genetic variability via RAPD of all collected isolates of *Pleurotus*.

## MATERIAL AND METHODS

### Sample collection

Total sixty isolates of genus *Pleurotus* were collected from the six identified geo-ecological zones of eastern Uttar Pradesh (India) namely: *usar*, wastelands, forest area, wetlands, flood area and fertile lands (Table 1). Thirty nine isolates could be preserved for further studies; rest were either much perished or sometimes highly contaminated. The collection of isolates was made as per the standard protocol for macro fungi outlined by Mueller et al. (2004), which were accompanied with recording of morphological feature such as size, color, texture and surface of pileus, nature of cuticle in lamelle on the pileus, position, size, texture and structure and size of stipe as given in table (Table 2). Remaining characteristics such as size and color of basidiospores were recorded in the laboratory. No two samples were collected within the distance of 4 feet since many fruiting bodies within a radius of few feet could be the outcome of the same mycelium. We have also recorded the radial growth of mycelia and shown in our previous study (Patel et al., 2017). Pure culture of the isolates were finally preserved on potato dextrose agar (PDA) slants and stored at 4°C, and were revived on fresh PDA slants at every 2-3 weeks interval.

**Table 1:** Details of geographic coordinates of studied districts with their geographical characters, in which one district could be comprises of more than one eco-edaphic zones.

| Geo-ecological zones | Districts | Altitude (m) | Latitude | Longitude |
| --- | --- | --- | --- | --- |
| **1. Usar** | Jaunpur | 79.5 - 88.3 | 24°24'N-26°12'N | 82°70'E-83°50'E |
|  | Allahabad | 72.0 – 98.0 | 24°87'N-25°27'N | 81°51'E-82°51'E |
|  | Bhadohi | 85.0 - 88.5 | 25°24'N-25°42'N | 82°38'E-82°57'E |
| **2. Foresting area** | Mirzapur | 80.0 - 128.9 | 23°32'N-25°52'N | 82°7'E-83°53'E |
| **3. Wetlands** | Allahabad | 72.0 – 98.0 | 24°87'N-25°27'N | 81°51'E-82°51'E |
|  | Varanasi | 75.7 - 80.7 | 25°14'N-25°23.5'N | 82°56'E-83° 8'E |
|  | Jaunpur | 79.5 - 88.3 | 24°24'N-26°12'N | 82°70'E-83°50'E |
|  | Mirzapur | 80.0 - 128.9 | 23°32'N-25°52'N | 82°7'E-83°53'E |
| **4. Flood area** | Jaunpur | 79.5 - 88.3 | 24°24'N-26°12'N | 82°70'E-83°50'E |
|  | Allahabad | 72.0 – 98.0 | 24°87'N-25°27'N | 81°51'E-82°51'E |
|  | Varanasi | 75.7 - 80.7 | 25°14'N-25°23.5'N | 82°56'E-83° 8'E |
|  | Azamgarh | 64.0 - 71.5 | 25°90'N-26°38'N | 82°40'E-83°53'E |
| **5. Fertile lands** | Allahabad | 72.0 – 98.0 | 24°87'N-25°27'N | 81°51'E-82°51'E |
|  | Azamgarh | 64.0 - 71.5 | 25°90'N-26°38'N | 82°40'E-83°53'E |
|  | Varanasi | 75.7 - 80.7 | 25°14'N-25°23.5'N | 82°56'E-83° 8'E |
|  | Jaunpur | 79.5 - 88.3 | 24°24'N-26°12'N | 82°70'E-83°50'E |
|  | Mirzapur | 80.0 - 128.9 | 23°32'N-25°52'N | 82°7'E-83°53'E |
| **6. Wastelands** | Mirzapur | 80.0 - 128.9 | 23°32'N-25°52'N | 82°7'E-83°53'E |
|  | Allahabad | 72.0 – 98.0 | 24°87'N-25°27'N | 81°51'E-82°51'E |
|  | Varanasi | 75.7 - 80.7 | 25°14'N-25°23.5'N | 82°56'E-83° 8'E |

**Table 2:**
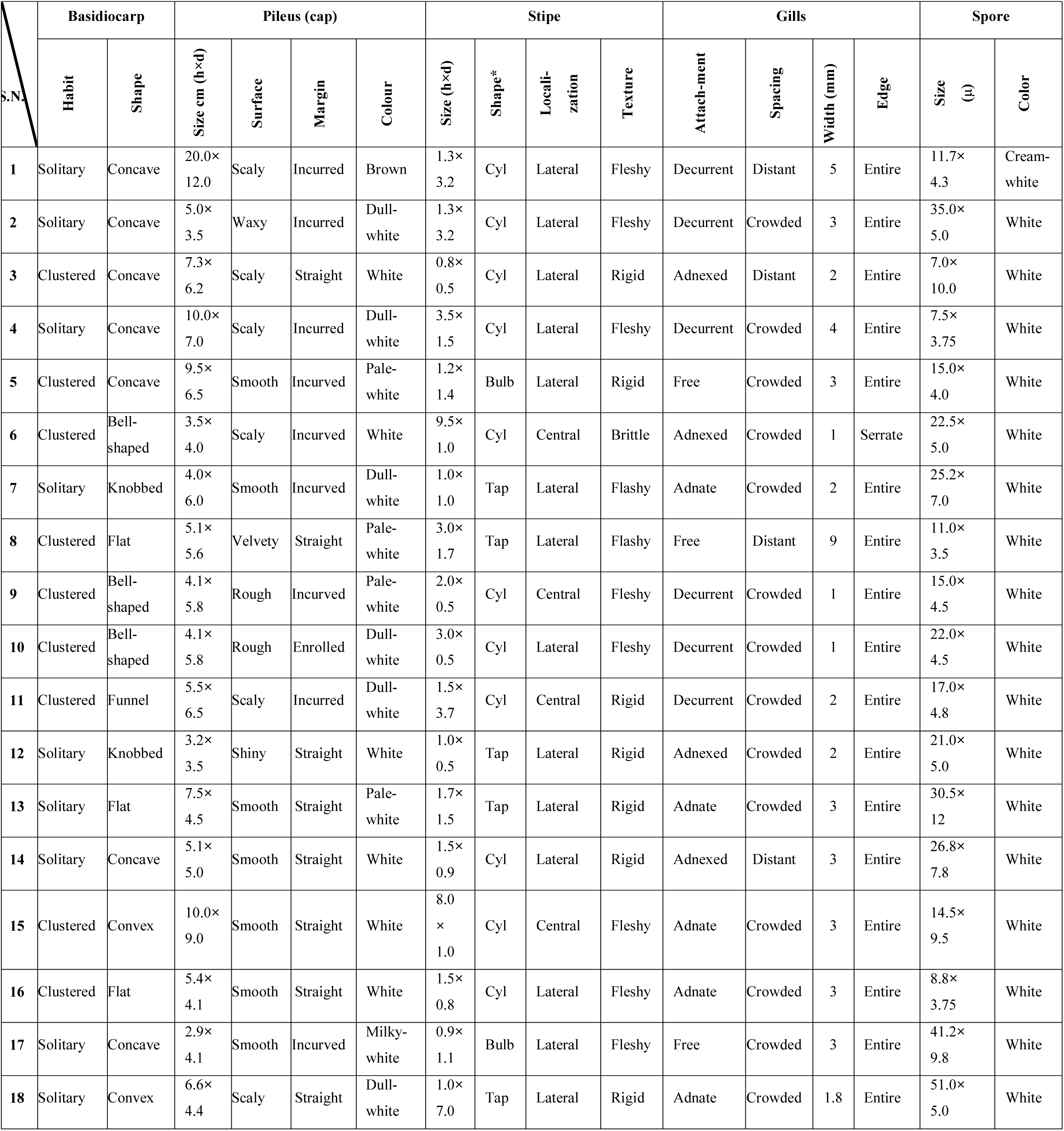
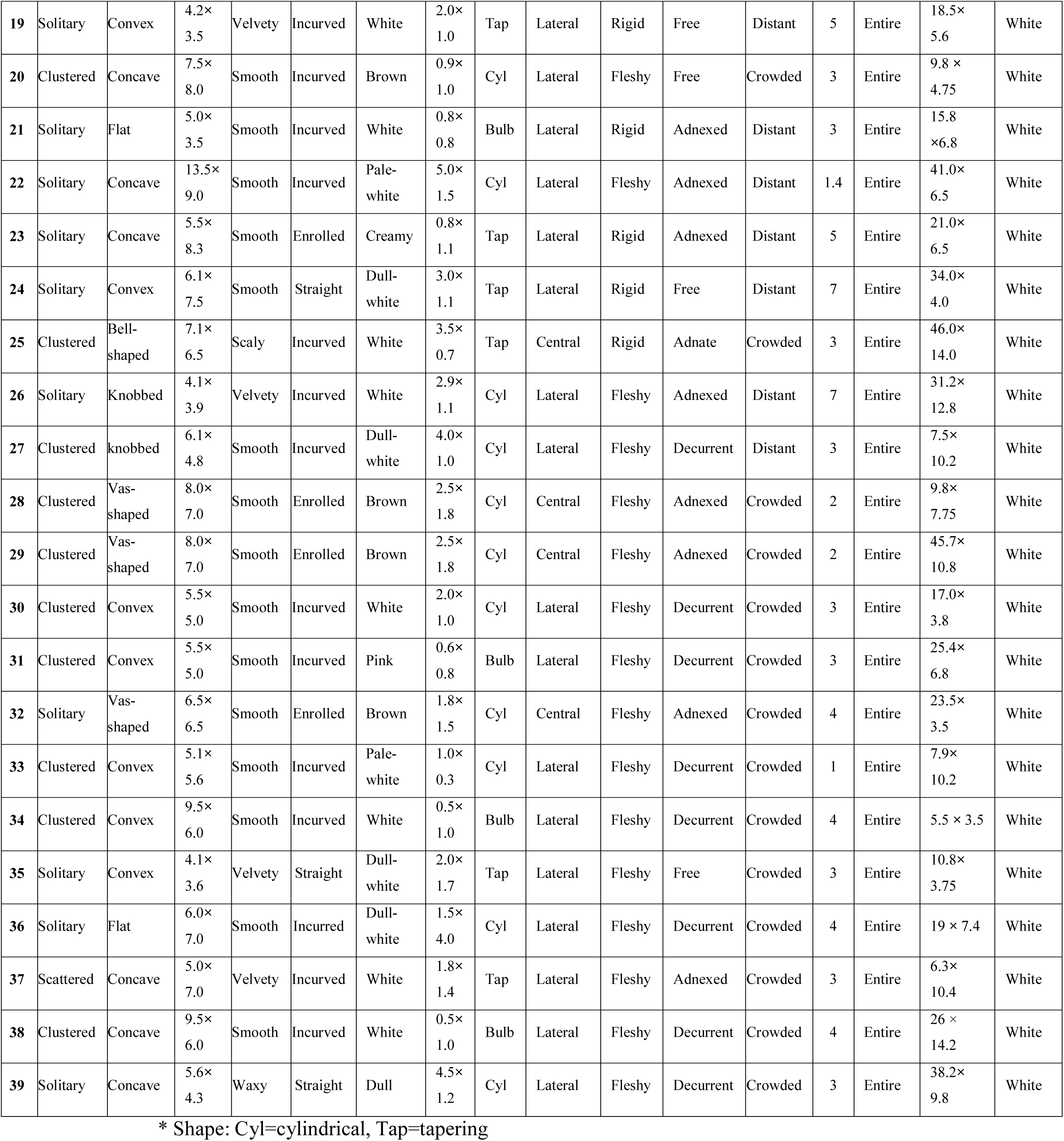
Morphological features of basidiocarps (fruiting bodies), their pileus, stipe, gills and spores of collected isolates of Pleurotus species

| S.N. | Basidiocarp |  | Pileus (cap) |  |  |  | Stipe |  |  |  | Gills |  |  |  | Spore |  |
| --- | --- | --- | --- | --- | --- | --- | --- | --- | --- | --- | --- | --- | --- | --- | --- | --- |
|  | Habit | Shape | Size cm (h×d) | Surface | Margin | Colour | Size (h×d) | Shape* | Locali-<br>zation | Texture | Attach-<br>ment | Spacing | Width (mm) | Edge | Size<br>(μ) | Color |
| 1 | Solitary | Concave | 20.0×<br>12.0 | Scaly | Incurved | Brown | 1.3×<br>3.2 | Cyl | Lateral | Fleshy | Decurrent | Distant | 5 | Entire | 11.7×<br>4.3 | Cream-<br>white |
| 2 | Solitary | Concave | 5.0×<br>3.5 | Waxy | Incurved | Dull-<br>white | 1.3×<br>3.2 | Cyl | Lateral | Fleshy | Decurrent | Crowded | 3 | Entire | 35.0×<br>5.0 | White |
| 3 | Clustered | Concave | 7.3×<br>6.2 | Scaly | Straight | White | 0.8×<br>0.5 | Cyl | Lateral | Rigid | Adnexed | Distant | 2 | Entire | 7.0×<br>10.0 | White |
| 4 | Solitary | Concave | 10.0×<br>7.0 | Scaly | Incurved | Dull-<br>white | 3.5×<br>1.5 | Cyl | Lateral | Fleshy | Decurrent | Crowded | 4 | Entire | 7.5×<br>3.75 | White |
| 5 | Clustered | Concave | 9.5×<br>6.5 | Smooth | Incurved | Pale-<br>white | 1.2×<br>1.4 | Bulb | Lateral | Rigid | Free | Crowded | 3 | Entire | 15.0×<br>4.0 | White |
| 6 | Clustered | Bell-<br>shaped | 3.5×<br>4.0 | Scaly | Incurved | White | 9.5×<br>1.0 | Cyl | Central | Brittle | Adnexed | Crowded | 1 | Serrate | 22.5×<br>5.0 | White |
| 7 | Solitary | Knobbed | 4.0×<br>6.0 | Smooth | Incurved | Dull-<br>white | 1.0×<br>1.0 | Tap | Lateral | Flashy | Adnate | Crowded | 2 | Entire | 25.2×<br>7.0 | White |
| 8 | Clustered | Flat | 5.1×<br>5.6 | Velvety | Straight | Pale-<br>white | 3.0×<br>1.7 | Tap | Lateral | Flashy | Free | Distant | 9 | Entire | 11.0×<br>3.5 | White |
| 9 | Clustered | Bell-<br>shaped | 4.1×<br>5.8 | Rough | Incurved | Pale-<br>white | 2.0×<br>0.5 | Cyl | Central | Fleshy | Decurrent | Crowded | 1 | Entire | 15.0×<br>4.5 | White |
| 10 | Clustered | Bell-<br>shaped | 4.1×<br>5.8 | Rough | Enrolled | Dull-<br>white | 3.0×<br>0.5 | Cyl | Lateral | Fleshy | Decurrent | Crowded | 1 | Entire | 22.0×<br>4.5 | White |
| 11 | Clustered | Funnel | 5.5×<br>6.5 | Scaly | Incurved | Dull-<br>white | 1.5×<br>3.7 | Cyl | Central | Rigid | Decurrent | Crowded | 2 | Entire | 17.0×<br>4.8 | White |
| 12 | Solitary | Knobbed | 3.2×<br>3.5 | Shiny | Straight | White | 1.0×<br>0.5 | Tap | Lateral | Rigid | Adnexed | Crowded | 2 | Entire | 21.0×<br>5.0 | White |
| 13 | Solitary | Flat | 7.5×<br>4.5 | Smooth | Straight | Pale-<br>white | 1.7×<br>1.5 | Tap | Lateral | Rigid | Adnate | Crowded | 3 | Entire | 30.5×<br>12 | White |
| 14 | Solitary | Concave | 5.1×<br>5.0 | Smooth | Straight | White | 1.5×<br>0.9 | Cyl | Lateral | Rigid | Adnexed | Distant | 3 | Entire | 26.8×<br>7.8 | White |
| 15 | Clustered | Convex | 10.0×<br>9.0 | Smooth | Straight | White | 8.0<br>×<br>1.0 | Cyl | Central | Fleshy | Adnate | Crowded | 3 | Entire | 14.5×<br>9.5 | White |
| 16 | Clustered | Flat | 5.4×<br>4.1 | Smooth | Straight | White | 1.5×<br>0.8 | Cyl | Lateral | Fleshy | Adnate | Crowded | 3 | Entire | 8.8×<br>3.75 | White |
| 17 | Solitary | Concave | 2.9×<br>4.1 | Smooth | Incurved | Milky-<br>white | 0.9×<br>1.1 | Bulb | Lateral | Fleshy | Free | Crowded | 3 | Entire | 41.2×<br>9.8 | White |
| 18 | Solitary | Convex | 6.6×<br>4.4 | Scaly | Straight | Dull-<br>white | 1.0×<br>7.0 | Tap | Lateral | Rigid | Adnate | Crowded | 1.8 | Entire | 51.0×<br>5.0 | White |
| 19 | Solitary | Convex | 4.2×<br>3.5 | Velvety | Incurved | White | 2.0×<br>1.0 | Tap | Lateral | Rigid | Free | Distant | 5 | Entire | 18.5×<br>5.6 | White |
| 20 | Clustered | Concave | 7.5×<br>8.0 | Smooth | Incurved | Brown | 0.9×<br>1.0 | Cyl | Lateral | Fleshy | Free | Crowded | 3 | Entire | 9.8 ×<br>4.75 | White |
| 21 | Solitary | Flat | 5.0×<br>3.5 | Smooth | Incurved | White | 0.8×<br>0.8 | Bulb | Lateral | Rigid | Adnexed | Distant | 3 | Entire | 15.8<br>×6.8 | White |
| 22 | Solitary | Concave | 13.5×<br>9.0 | Smooth | Incurved | Pale-<br>white | 5.0×<br>1.5 | Cyl | Lateral | Fleshy | Adnexed | Distant | 1.4 | Entire | 41.0×<br>6.5 | White |
| 23 | Solitary | Concave | 5.5×<br>8.3 | Smooth | Enrolled | Creamy | 0.8×<br>1.1 | Tap | Lateral | Rigid | Adnexed | Distant | 5 | Entire | 21.0×<br>6.5 | White |
| 24 | Solitary | Convex | 6.1×<br>7.5 | Smooth | Straight | Dull-<br>white | 3.0×<br>1.1 | Tap | Lateral | Rigid | Free | Distant | 7 | Entire | 34.0×<br>4.0 | White |
| 25 | Clustered | Bell-<br>shaped | 7.1×<br>6.5 | Scaly | Incurved | White | 3.5×<br>0.7 | Tap | Central | Rigid | Adnate | Crowded | 3 | Entire | 46.0×<br>14.0 | White |
| 26 | Solitary | Knobbed | 4.1×<br>3.9 | Velvety | Incurved | White | 2.9×<br>1.1 | Cyl | Lateral | Fleshy | Adnexed | Distant | 7 | Entire | 31.2×<br>12.8 | White |
| 27 | Clustered | knobbed | 6.1×<br>4.8 | Smooth | Incurved | Dull-<br>white | 4.0×<br>1.0 | Cyl | Lateral | Fleshy | Decurrent | Distant | 3 | Entire | 7.5×<br>10.2 | White |
| 28 | Clustered | Vas-shaped | 8.0×<br>7.0 | Smooth | Enrolled | Brown | 2.5×<br>1.8 | Cyl | Central | Fleshy | Adnexed | Crowded | 2 | Entire | 9.8×<br>7.75 | White |
| 29 | Clustered | Vas-shaped | 8.0×<br>7.0 | Smooth | Enrolled | Brown | 2.5×<br>1.8 | Cyl | Central | Fleshy | Adnexed | Crowded | 2 | Entire | 45.7×<br>10.8 | White |
| 30 | Clustered | Convex | 5.5×<br>5.0 | Smooth | Incurved | White | 2.0×<br>1.0 | Cyl | Lateral | Fleshy | Decurrent | Crowded | 3 | Entire | 17.0×<br>3.8 | White |
| 31 | Clustered | Convex | 5.5×<br>5.0 | Smooth | Incurved | Pink | 0.6×<br>0.8 | Bulb | Lateral | Fleshy | Decurrent | Crowded | 3 | Entire | 25.4×<br>6.8 | White |
| 32 | Solitary | Vas-shaped | 6.5×<br>6.5 | Smooth | Enrolled | Brown | 1.8×<br>1.5 | Cyl | Central | Fleshy | Adnexed | Crowded | 4 | Entire | 23.5×<br>3.5 | White |
| 33 | Clustered | Convex | 5.1×<br>5.6 | Smooth | Incurved | Pale-<br>white | 1.0×<br>0.3 | Cyl | Lateral | Fleshy | Decurrent | Crowded | 1 | Entire | 7.9×<br>10.2 | White |
| 34 | Clustered | Convex | 9.5×<br>6.0 | Smooth | Incurved | White | 0.5×<br>1.0 | Bulb | Lateral | Fleshy | Decurrent | Crowded | 4 | Entire | 5.5 × 3.5 | White |
| 35 | Solitary | Convex | 4.1×<br>3.6 | Velvety | Straight | Dull-<br>white | 2.0×<br>1.7 | Tap | Lateral | Fleshy | Free | Crowded | 3 | Entire | 10.8×<br>3.75 | White |
| 36 | Solitary | Flat | 6.0×<br>7.0 | Smooth | Incurved | Dull-<br>white | 1.5×<br>4.0 | Cyl | Lateral | Fleshy | Decurrent | Crowded | 4 | Entire | 19 × 7.4 | White |
| 37 | Scattered | Concave | 5.0×<br>7.0 | Velvety | Incurved | White | 1.8×<br>1.4 | Tap | Lateral | Fleshy | Adnexed | Crowded | 3 | Entire | 6.3×<br>10.4 | White |
| 38 | Clustered | Concave | 9.5×<br>6.0 | Smooth | Incurved | White | 0.5×<br>1.0 | Bulb | Lateral | Fleshy | Decurrent | Crowded | 4 | Entire | 26 ×<br>14.2 | White |
| 39 | Solitary | Concave | 5.6×<br>4.3 | Waxy | Straight | Dull | 4.5×<br>1.2 | Cyl | Lateral | Fleshy | Decurrent | Crowded | 3 | Entire | 38.2×<br>9.8 | White |

\* Shape: Cyl=cylindrical, Tap=tapering

### Characterization of isolates through PCR-fingerprinting

For molecular characterization of isolates through RAPD, genomic DNA was isolated from mycelia of pure cultures grown in potato dextrose broth by CTAB method as described by Sadowsky et al. (1987) and Kuramae-Izioka (1997). Only high quality DNA (260/280 = 1.7-1.9) were used in this study. Genetic diversity within collected isolates was characterized by PCR using arbitrary ten-mer primers as given in table (Table 3). Amplification was carried out according to the methods of Williams et al. (1990) in biological triplicate. PCR reactions were performed in thermal cycler (Bio-Rad). The reaction volume (25 μl) contained DNA (40–100 ng), *Taq* buffer including 1.5 mM MgCl_2_, 10mM dNTPs, 10 pM of a primer and 1-2 units of *Taq* DNA polymerase with negative control without template. Amplification reaction included i) initial denaturation at 95°C for 5 min, ii) 35 cycles of denaturation at 94°C for 60 sec, primer annealing at ∼32 °C (varied with primers) for 30 sec and DNA synthesis at 72°C for 150 sec; and iii) final amplification at 72°C for 10 min. Amplified products were resolved on 1.2% agarose gel along with 100 bp DNA ladder. The gels were observed under Gel-Doc system (Alfa Imager) and pictured images used for further scoring of the banding patterns in different isolates.

**Table 3:**
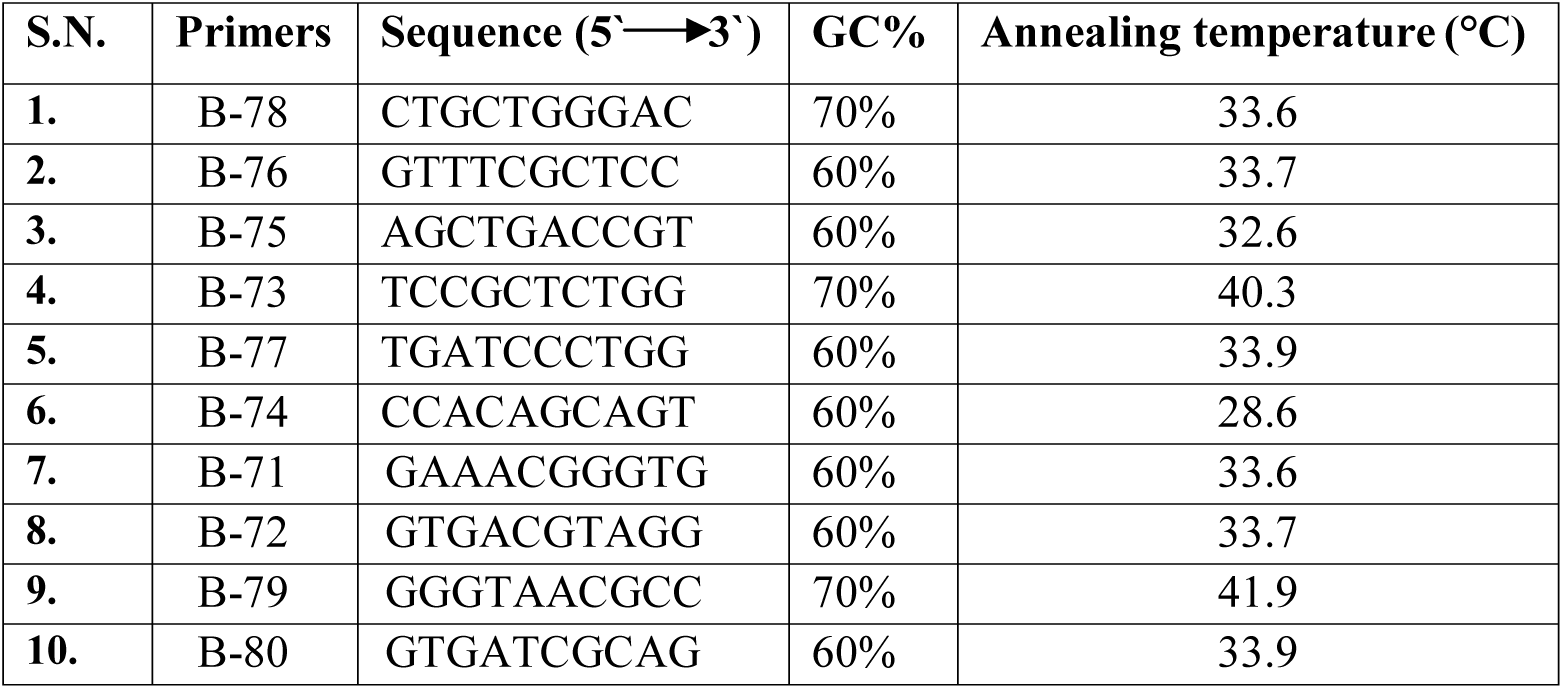
Primers with their sequence, GC% and Annealing temperature

### Quantification of nutraceuticals (proteins, carbohydrates, phenolics)

The total protein content present in dried fruiting bodies was analyzed by the standard method of Bradford (Bradford, 1976) and total carbohydrate content available in mycelium was determined using the phenol-sulfuric acid method of Dubois et al. (1956), while total amount of phenolics content of methanolic extract of dried mycelia was measured as per standard method developed by Singleton et al. (1965).

### Bioassay of Vitamin B_12_

Bioassay of vitamin B_12_ was conducted based upon the methods of AOAC (Williams, 2005). In which, Vitamin B_12_ assay medium (minimal medium) contains all other vitamins/nutrients essential for the growth of test organism (*Lactobacillus delbrueckii* subsp. *lactis*) except vitamin B_12_. The assay medium was prepared in double strength and 5 ml medium was taken in each test tube (Made: Borosil) to which increasing amounts of standard solution of or the unknown & sufficient water was added to give a total volume of 10 ml per tube. All the tubes were sterilized for 5 min at 10 psi and immediately cooled at room temperature. 100 µl of the inoculums (fresh culture of *Lactobacillus delbrueckii* subsp. *Lactis*) were inoculated into each of the assay tubes. Growth of bacteria in assay medium was measured at 530 nm by spectrophotometer (1 cm path length) after incubation for 36 h at 37 ±1 °C in shaker.

### Evaluation of laccase activity

Laccase activity was determined via the oxidation of Guaiacol (o-methoxyphenol catechol monomethylether) as substrate as per the method used by Arora and Sandhu (1985) and details given by Patel et al. (2015). The enzyme extracts were prepared by homogenization of mycelial mat in buffer in cold condition and then the activity was calculated as per the formula: IU/ml= <u>ΔA@470nm/0.001</u>

### Dye decolorization

To investigate the decolorization potential of the isolates, two most used dyes in textile industries; Bromophenol Blue (BPB) and Malachite greenG (MG) were procured from Himedia BioSciences (India) and were evaluated on solid medium through all the collected isolates. It was based on the measurement of bleached area by mycelia growth on PDA solid medium either supplemented with test dyes or without dye as control. Screening test for degradation of BPB and MG was carried out according to the method of Machado et al. (2005). For decolorization assay, PDA plates were prepared with 0.01% (w/v) MG supplementation. Point inoculation was performed on both types of plate as mycelial plugs prepared from pure mycelial mother plate with the help of cork-borer and kept at the center of the PDA plates; plates were incubated at 28°C±1 in BOD incubator. Similarly, 0.05% (w/v) BPB was supplemented with PDA. The decolorization of dyes was evaluated by measuring the clear zone formation under and around the developing mycelia from center of the PDA plates (plate diameter: 90.0 mm). Results were observed after 5 and 10 days, clear zone was appeared against blue background. The experiments were performed in biological triplicates.

### Statistical analysis

Analysis of mean, SD and SE were calculated by Microsoft excel. However, analysis of variability and allele frequency, allele number, effective allele number, polymorphic loci, observed homozygosity, expected homozygosity, Shannon Index (Gillies, 1997), Gene diversity, neutrality test (Manly, 1985) [32] and unbiased genetic distance were calculated by POPGENE 32 software. Pair-wise genetic dissimilarities among the isolates were calculated from the binary data using well known Jaccard’s Coefficient to form matrix of genetic dissimilarities. Dendrogram was obtained by using NTSYS-PC software version 2.02j through the RAPD binary matrix from which cluster analysis was performed by means of unweighted pair group method using arithmetic average (UPGMA). The variability among the isolates was assessed by comparing RAPD fragments according to their sizes and the presence/absence of shared fragments. All the statistical analyses related to nutritional elements were conducted using the SPAR v. 2.0.

## Results

### Collection and purification of isolates

Eco-edaphic zones identified for the present study as given in table (Table 1) and depicted in figure (Fig. 1) were thoroughly observed and identified in rainy season for hotspot of *Pleurotus* mushroom. Isolates were collected from the habituated dead and decayed mango (*Mangifera indica* Linn.) trees. A total of 39 isolates were purified out of 60 collected isolates from different zones and their morphological features (Table 2) were observed during and after collection. Interestingly, it was observed from the table (Table 2) that mixed types of *Pleurotus* were collected from different zones; which indicates that studied areas are diversity rich. Fruiting bodies were found both clustered as well as solitary types, however, shape of basidiocarps were diverse ranging from concave, bell shaped, funnel shaped, convex, knobbed, Vas-shaped to flat having scaly, waxy, velvety, rough or even smooth surface with varying in area ranging from 8.12 cm^2^ to 240 cm^2^. Fruiting bodies of collected isolates were of various colors such as brown, white, milky white, pink; which demonstrated the importance of study of diversity through the collection of natural isolates for further strain improvements and betterment of various potentialities. Stipes of isolates were found very interesting having majority in fleshy nature, however, few of them were with rigid stipe. Majority of stipes were cylindrical in shape, some of them were in tapering and few were in bulb shape. There was also diversity in attachment of stipes from its pileus, majority of them were in lateral, which is a signature characteristics of Oyster mushrooms (*Pleurotus* spp.), and however, few of them were centrally attached with its pileus. Gills found under the pileus were majorly in crowded with few distantly located having entire completed edge, some of them were serrated. Spores collected from fruiting bodies cultivated in lab were white in color and varying in size.

**Figure 1:**
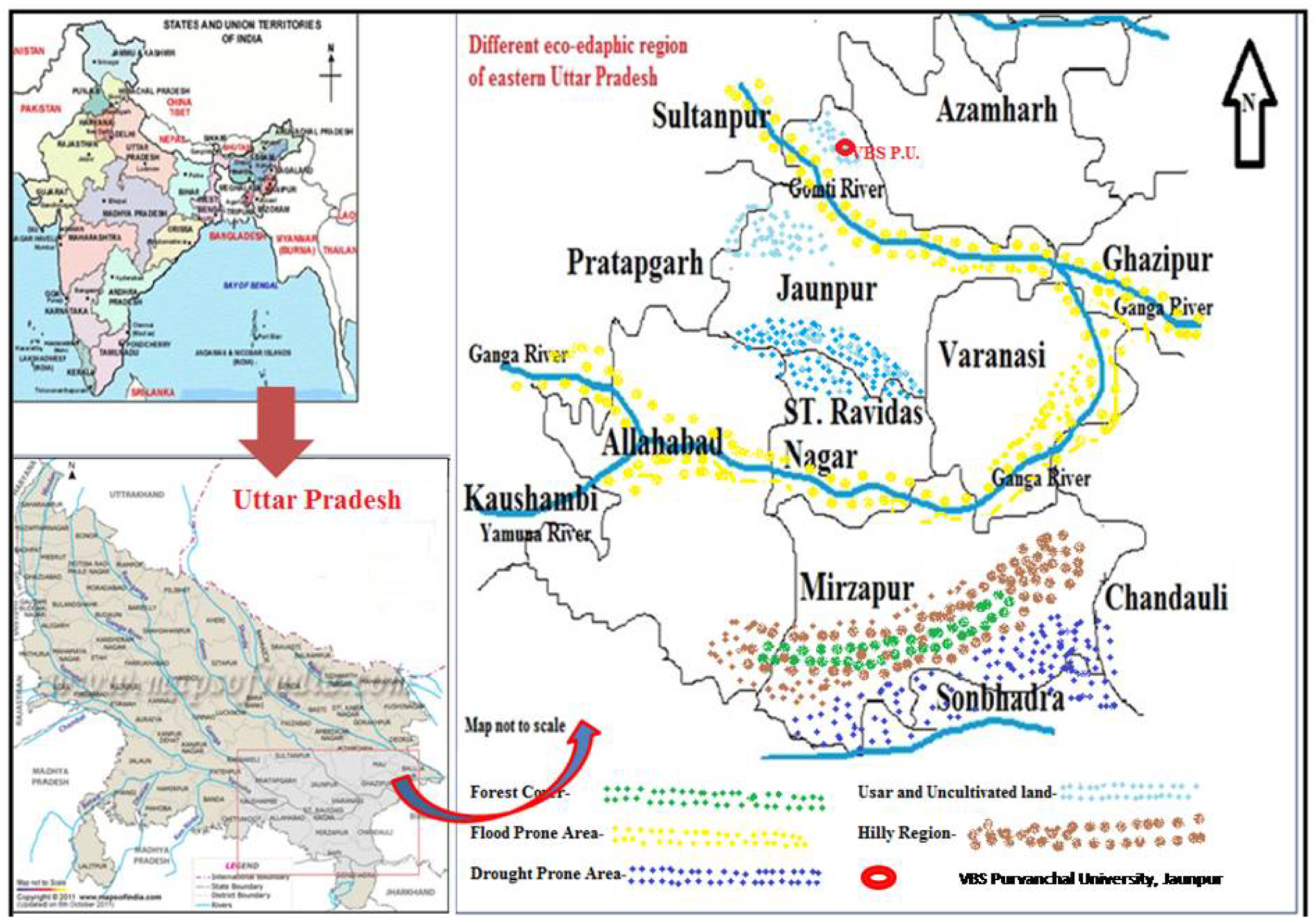
Selected zone in studied region. Eco-edaphic zones of selected area under investigation for the study of genetic diversity of *Pleurotus* species. Enlarge map of selected area from the State depicted right side of the figure. Dots in different color represent five different zone and rest white background represents fertile land. Arrow in the upper right most corner indicates direction north. There is two major rivers (Ganga and Yamuna) and many small rivers (Varuna, Gomti, Sai, etc) in the studied zones.

### DNA polymorphism through RAPD profiling and their allelic frequency

For the study of DNA polymorphism, genomic DNA from all the isolates was subjected to RAPD based DNA profiling to detect variability in the genome. Ten single strand primers (arbitrary primers) were chosen for amplification. The sizes of amplification products varied from 100-2500 bp and the degree of polymorphism depended upon the isolate and the primer employed. For convenience the results of RAPD profiling was broadly categorized into three groups as given in table 4 depending upon the degree of polymorphism as evident from the banding pattern of amplified products on agarose gels. The banding pattern of representative primer belonging to each of these groups as separated on agarose gel after PCR amplification (Gels are not shown here). Out of 10 selected primers used in this study, 6 of them amplified the DNA and generated 51 polymorphic bands given in table 4. The results clearly indicate that primers-B-73, B-74, B-75, B-76, B-77 and B-78 produced 8, 6, 9, 10, 6, and 12, respectively discrete and scorable bands in all isolates as given in table 4. Primer B-78 amplified the highest number of scorable bands (i.e. 12), whereas, primer B-74 and B-77 amplified the lowermost number of bands (i.e. 6). Each sample was characterized by a different RAPD genotype. For the assessment of genetic diversity of isolates, the frequencies of all resolved RAPD alleles (e.g. from B-73, B-74, B-75, B-76, B-77 and B-78) were calculated as given in table 5. The allele 1 having highest frequency has great importance in diversity study.

**Table 4:**
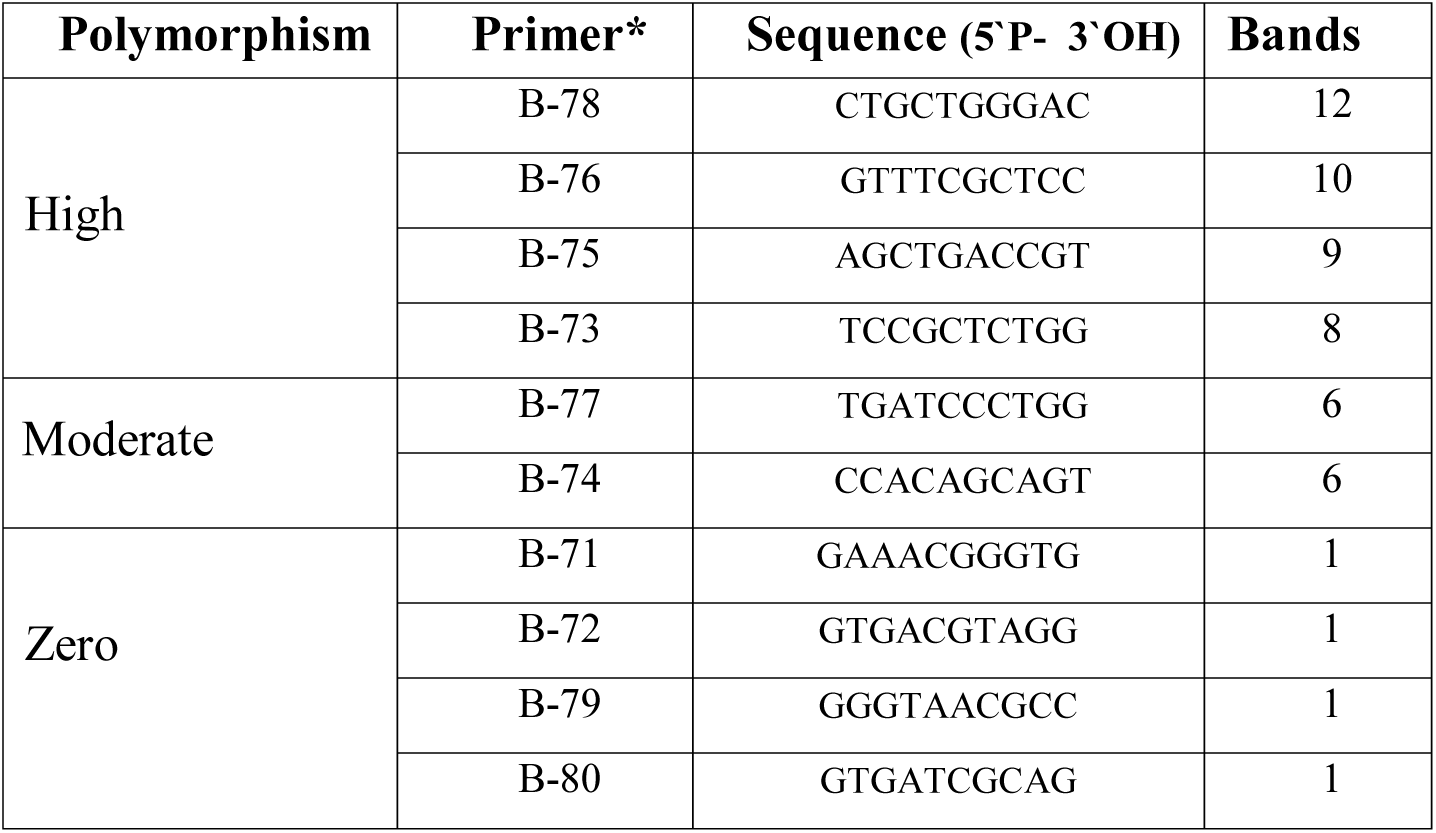
Different primers and their polymorphism status

**Table 5:** Frequency of genes/alleles in isolates

| Locus | Alleles |  | Locus | Alleles |  |
| --- | --- | --- | --- | --- | --- |
|  | Allele 1 | Allele 0 |  | Allele 1 | Allele 0 |
| *B73-1* | 0.152 | 0.847 | *B76-4* | 0.302 | 0.698 |
| *B73-2* | 0.122 | 0.877 | *B76-5* | 0.215 | 0.784 |
| *B73-3* | 0.152 | 0.847 | *B76-6* | 0.152 | 0.847 |
| *B73-4* | 0.094 | 0.905 | *B76-7* | 0.232 | 0.767 |
| *B73-5* | 0.167 | 0.832 | *B76-8* | 0.232 | 0.767 |
| *B73-6* | 0.152 | 0.847 | *B76-9* | 0.094 | 0.905 |
| *B73-7* | 0.026 | 0.974 | *B76-10* | 0.012 | 0.987 |
| *B73-8* | 0.137 | 0.862 | *B77-1* | 0.232 | 0.767 |
| *B74-1* | 0.066 | 0.933 | *B77-2* | 0.108 | 0.891 |
| *B74-2* | 0.215 | 0.784 | *B77-3* | 0.039 | 0.960 |
| *B74-3* | 0.122 | 0.877 | *B77-4* | 0.199 | 0.800 |
| *B74-4* | 0.167 | 0.832 | *B77-5* | 0.052 | 0.947 |
| *B74-5* | 0.094 | 0.905 | *B77-6* | 0.012 | 0.987 |
| *B74-6* | 0.094 | 0.905 | *B78-1* | 0.026 | 0.974 |
| *B75-1* | 0.094 | 0.905 | *B78-2* | 0.137 | 0.862 |
| *B75-2* | 0.232 | 0.767 | *B78-3* | 0.152 | 0.847 |
| *B75-3* | 0.248 | 0.751 | *B78-4* | 0.199 | 0.800 |
| *B75-4* | 0.167 | 0.832 | *B78-5* | 0.266 | 0.733 |
| *B75-5* | 0.137 | 0.862 | *B78-6* | 0.183 | 0.816 |
| *B75-6* | 0.283 | 0.716 | *B78-7* | 0.152 | 0.847 |
| *B75-7* | 0.400 | 0.599 | *B78-8* | 0.183 | 0.816 |
| *B75-8* | 0.167 | 0.832 | *B78-9* | 0.183 | 0.816 |
| *B75-9* | 0.066 | 0.933 | *B78-10* | 0.039 | 0.960 |
| *B76-1* | 0.012 | 0.987 | *B78-11* | 0.122 | 0.877 |
| *B76-2* | 0.422 | 0.577 | *B78-12* | 0.108 | 0.891 |
| *B76-3* | 0.199 | 0.800 |  |  |  |

### Genetic diversity among isolates

Many descriptive measures of diversity are calculated for the estimation of diversity among collected isolates from different ecological zones as described by Nei (1973). These descriptive measures are: number of loci analyzed, percent polymorphic loci, mean number of alleles per locus, gene diversity, effective number of alleles (Kimura and Crow, 1964), observed heterozygosity and Shannon’s Information Index (Lewontin, 1972) as given in table (Table 6). The isolates were assigned to 39 different subtypes by RAPD analysis. The average gene diversity (as per Nei’s) for all RAPD loci was 0.244±0.116. The minimum gene diversity was shown by *B76*-1 loci, i.e. 0.025 while maximum diversity was shown by *B76*-2 loci, i.e. 0.488. The observed number of alleles was 2.000±0.000 while effective number of alleles was 1.355±0.215 (Table 6). Shannon’s Information Index was found to 0.397±0.155. The Nei’s (1978) unbiased genetic similarity among all pairs of samples is given in table 7. This similarity matrix clearly showed the minimum 11%, and maximum 84% similarity between isolates #7 and #8. The Ewens-Watterson test for neutrality for each locus was calculated by POPGENE as given in table 8 showed that the allele frequencies at all loci were selectively neutral in the studied isolates. Where for each observed allele frequency, upper (U95) and lower (L95) were at 95% confidence limits of expected F values.

**Table 6:** Summary of gene diversity for all RAPD’s loci as per Nei (1973)

| Locus | Sample Size | Number of alleles |  | Gene diversity (h) | Shannon's index (I) |
| --- | --- | --- | --- | --- | --- |
|  |  | Observed (na) | Effective (ne) |  |  |
| B73-1 | 39 | 2.0000 | 1.3491 | 0.2587 | 0.4273 |
| B73-2 | 39 | 2.0000 | 1.2749 | 0.2157 | 0.3727 |
| B73-3 | 39 | 2.0000 | 1.3491 | 0.2587 | 0.4273 |
| B73-4 | 39 | 2.0000 | 1.2057 | 0.1706 | 0.3121 |
| B73-5 | 39 | 2.0000 | 1.3879 | 0.2795 | 0.4526 |
| B73-6 | 39 | 2.0000 | 1.3491 | 0.2587 | 0.4273 |
| B73-7 | 39 | 2.0000 | 1.0533 | 0.0506 | 0.1205 |
| B73-8 | 39 | 2.0000 | 1.3114 | 0.2375 | 0.4007 |
| B74-1 | 39 | 2.0000 | 1.1413 | 0.1238 | 0.2440 |
| B74-2 | 39 | 2.0000 | 1.5109 | 0.3382 | 0.5212 |
| B74-3 | 39 | 2.0000 | 1.2749 | 0.2157 | 0.3727 |
| B74-4 | 39 | 2.0000 | 1.3879 | 0.2795 | 0.4526 |
| B74-5 | 39 | 2.0000 | 1.2057 | 0.1706 | 0.3121 |
| B74-6 | 39 | 2.0000 | 1.2057 | 0.1706 | 0.3121 |
| B75-1 | 39 | 2.0000 | 1.2057 | 0.1706 | 0.3121 |
| B75-2 | 39 | 2.0000 | 1.5538 | 0.3564 | 0.5417 |
| B75-3 | 39 | 2.0000 | 1.5973 | 0.3739 | 0.5612 |
| B75-4 | 39 | 2.0000 | 1.3879 | 0.2795 | 0.4526 |
| B75-5 | 39 | 2.0000 | 1.3114 | 0.2375 | 0.4007 |
| B75-6 | 39 | 2.0000 | 1.6852 | 0.4066 | 0.5966 |
| B75-7 | 39 | 2.0000 | 1.9243 | 0.4803 | 0.6734 |
| B75-8 | 39 | 2.0000 | 1.3879 | 0.2795 | 0.4526 |
| B75-9 | 39 | 2.0000 | 1.1413 | 0.1238 | 0.2440 |
| B76-1 | 39 | 2.0000 | 1.0261 | 0.0255 | 0.0690 |
| B76-2 | 39 | 2.0000 | 1.9533 | 0.4880 | 0.6811 |
| B76-3 | 39 | 2.0000 | 1.4689 | 0.3192 | 0.4995 |
| B76-4 | 39 | 2.0000 | 1.7289 | 0.4216 | 0.6126 |
| B76-5 | 39 | 2.0000 | 1.5109 | 0.3382 | 0.5212 |
| B76-6 | 39 | 2.0000 | 1.3491 | 0.2587 | 0.4273 |
| B76-7 | 39 | 2.0000 | 1.5538 | 0.3564 | 0.5417 |
| B76-8 | 39 | 2.0000 | 1.5538 | 0.3564 | 0.5417 |
| B76-9 | 39 | 2.0000 | 1.2057 | 0.1706 | 0.3121 |
| B76-10 | 39 | 2.0000 | 1.0261 | 0.0255 | 0.0690 |
| B77-1 | 39 | 2.0000 | 1.5538 | 0.3564 | 0.5417 |
| B77-2 | 39 | 2.0000 | 1.2397 | 0.1934 | 0.3433 |
| B77-3 | 39 | 2.0000 | 1.0815 | 0.0754 | 0.1655 |
| B77-4 | 39 | 2.0000 | 1.4689 | 0.3192 | 0.4995 |
| B77-5 | 39 | 2.0000 | 1.1109 | 0.0998 | 0.2063 |
| B77-6 | 39 | 2.0000 | 1.0261 | 0.0255 | 0.0690 |
| B78-1 | 39 | 2.0000 | 1.0533 | 0.0506 | 0.1205 |
| B78-2 | 39 | 2.0000 | 1.3114 | 0.2375 | 0.4007 |
| B78-3 | 39 | 2.0000 | 1.3491 | 0.2587 | 0.4273 |
| B78-4 | 39 | 2.0000 | 1.4689 | 0.3192 | 0.4995 |
| B78-5 | 39 | 2.0000 | 1.6412 | 0.3907 | 0.5794 |
| B78-6 | 39 | 2.0000 | 1.4279 | 0.2997 | 0.4767 |
| B78-7 | 39 | 2.0000 | 1.3491 | 0.2587 | 0.4273 |
| B78-8 | 39 | 2.0000 | 1.4279 | 0.2997 | 0.4767 |
| B78-9 | 39 | 2.0000 | 1.4279 | 0.2997 | 0.4767 |
| B78-10 | 39 | 2.0000 | 1.0815 | 0.0754 | 0.1655 |
| B78-11 | 39 | 2.0000 | 1.2749 | 0.2157 | 0.3727 |
| B78-12 | 39 | 2.0000 | 1.2397 | 0.1934 | 0.3433 |
| **Mean** | **39** | **2.0000** | **1.3551** | **0.2444** | **0.3972** |
| **SD Dev** |  | **0.0000** | **0.2148** | **0.1159** | **0.1547** |

**Table 7:** Nei’s unbiased measures of genetic similarity (Nei, 1978)

|  | **1** | **2** | **3** | **4** | **5** | **6** | **7** | **8** | **9** | **10** | **11** | **12** | **13** |
| --- | --- | --- | --- | --- | --- | --- | --- | --- | --- | --- | --- | --- | --- |
| **1** | 1.00 |  |  |  |  |  |  |  |  |  |  |  |  |
| **2** | 0.45 | 1.00 |  |  |  |  |  |  |  |  |  |  |  |
| **3** | 0.20 | 0.25 | 1.00 |  |  |  |  |  |  |  |  |  |  |
| **4** | 0.45 | 0.63 | 0.30 | 1.00 |  |  |  |  |  |  |  |  |  |
| **5** | 0.20 | 0.30 | 0.45 | 0.30 | 1.00 |  |  |  |  |  |  |  |  |
| **6** | 0.14 | 0.27 | 0.18 | 0.21 | 0.18 | 1.00 |  |  |  |  |  |  |  |
| **7** | 0.52 | 0.63 | 0.25 | 0.63 | 0.17 | 0.16 | 1.00 |  |  |  |  |  |  |
| **8** | 0.55 | 0.59 | 0.22 | 0.66 | 0.22 | 0.12 | 0.84 | 1.00 |  |  |  |  |  |
| **9** | 0.16 | 0.14 | 0.36 | 0.14 | 0.15 | 0.04 | 0.18 | 0.14 | 1.00 |  |  |  |  |
| **10** | 0.23 | 0.38 | 0.54 | 0.44 | 0.41 | 0.33 | 0.33 | 0.29 | 0.28 | 1.00 |  |  |  |
| **11** | 0.27 | 0.33 | 0.25 | 0.33 | 0.30 | 0.20 | 0.28 | 0.29 | 0.07 | 0.33 | 1.00 |  |  |
| **12** | 0.22 | 0.27 | 0.52 | 0.42 | 0.34 | 0.11 | 0.32 | 0.28 | 0.43 | 0.48 | 0.26 | 1.00 |  |
| **13** | 0.20 | 0.21 | 0.39 | 0.30 | 0.45 | 0.13 | 0.25 | 0.17 | 0.25 | 0.30 | 0.30 | 0.45 | 1.00 |
| **14** | 0.23 | 0.24 | 0.38 | 0.19 | 0.38 | 0.21 | 0.14 | 0.15 | 0.35 | 0.34 | 0.23 | 0.28 | 0.26 |
| **15** | 0.27 | 0.23 | 0.30 | 0.18 | 0.30 | 0.04 | 0.23 | 0.19 | 0.27 | 0.14 | 0.33 | 0.26 | 0.42 |
| **16** | 0.17 | 0.10 | 0.31 | 0.14 | 0.26 | 0.00 | 0.19 | 0.15 | 0.42 | 0.29 | 0.23 | 0.33 | 0.38 |
| **17** | 0.23 | 0.20 | 0.30 | 0.28 | 0.25 | 0.07 | 0.28 | 0.25 | 0.39 | 0.24 | 0.18 | 0.37 | 0.30 |
| **18** | 0.17 | 0.24 | 0.20 | 0.19 | 0.31 | 0.04 | 0.19 | 0.20 | 0.23 | 0.19 | 0.35 | 0.33 | 0.38 |
| **19** | 0.27 | 0.18 | 0.25 | 0.23 | 0.15 | 0.09 | 0.28 | 0.24 | 0.27 | 0.23 | 0.16 | 0.22 | 0.11 |
| **20** | 0.16 | 0.26 | 0.19 | 0.22 | 0.10 | 0.08 | 0.32 | 0.28 | 0.16 | 0.17 | 0.11 | 0.17 | 0.03 |
| **21** | 0.21 | 0.18 | 0.25 | 0.18 | 0.15 | 0.04 | 0.33 | 0.24 | 0.21 | 0.18 | 0.12 | 0.17 | 0.15 |
| **22** | 0.10 | 0.18 | 0.14 | 0.08 | 0.20 | 0.20 | 0.04 | 0.04 | 0.10 | 0.18 | 0.22 | 0.12 | 0.09 |
| **23** | 0.08 | 0.20 | 0.21 | 0.20 | 0.21 | 0.15 | 0.11 | 0.11 | 0.18 | 0.20 | 0.18 | 0.24 | 0.16 |
| **24** | 0.13 | 0.30 | 0.16 | 0.20 | 0.16 | 0.17 | 0.15 | 0.16 | 0.18 | 0.11 | 0.18 | 0.19 | 0.07 |
| **25** | 0.17 | 0.24 | 0.16 | 0.19 | 0.16 | 0.09 | 0.19 | 0.20 | 0.12 | 0.19 | 0.23 | 0.14 | 0.11 |
| **26** | 0.09 | 0.12 | 0.18 | 0.03 | 0.18 | 0.06 | 0.07 | 0.08 | 0.14 | 0.03 | 0.14 | 0.16 | 0.13 |
| **27** | 0.14 | 0.23 | 0.28 | 0.15 | 0.28 | 0.11 | 0.18 | 0.12 | 0.36 | 0.25 | 0.17 | 0.25 | 0.28 |
| **28** | 0.12 | 0.19 | 0.12 | 0.19 | 0.26 | 0.09 | 0.19 | 0.20 | 0.12 | 0.14 | 0.23 | 0.14 | 0.26 |
| **29** | 0.23 | 0.24 | 0.25 | 0.16 | 0.17 | 0.03 | 0.24 | 0.20 | 0.33 | 0.24 | 0.18 | 0.27 | 0.17 |
| **30** | 0.12 | 0.24 | 0.21 | 0.10 | 0.45 | 0.09 | 0.14 | 0.15 | 0.12 | 0.19 | 0.28 | 0.23 | 0.31 |
| **31** | 0.23 | 0.14 | 0.15 | 0.18 | 0.25 | 0.15 | 0.17 | 0.11 | 0.12 | 0.10 | 0.28 | 0.10 | 0.16 |
| **32** | 0.12 | 0.14 | 0.15 | 0.10 | 0.15 | 0.09 | 0.14 | 0.14 | 0.21 | 0.06 | 0.21 | 0.13 | 0.20 |
| **33** | 0.13 | 0.07 | 0.08 | 0.07 | 0.17 | 0.10 | 0.07 | 0.07 | 0.08 | 0.07 | 0.25 | 0.07 | 0.17 |
| **34** | 0.04 | 0.03 | 0.23 | 0.00 | 0.18 | 0.11 | 0.00 | 0.00 | 0.14 | 0.12 | 0.14 | 0.16 | 0.08 |
| **35** | 0.21 | 0.12 | 0.13 | 0.08 | 0.08 | 0.00 | 0.17 | 0.13 | 0.15 | 0.08 | 0.15 | 0.07 | 0.08 |
| **36** | 0.15 | 0.17 | 0.13 | 0.17 | 0.19 | 0.00 | 0.23 | 0.18 | 0.04 | 0.12 | 0.27 | 0.21 | 0.13 |
| **37** | 0.08 | 0.15 | 0.16 | 0.11 | 0.12 | 0.04 | 0.20 | 0.1 | 0.08 | 0.11 | 0.18 | 0.19 | 0.16 |
| **38** | 0.20 | 0.17 | 0.29 | 0.13 | 0.14 | 0.25 | 0.17 | 0.10 | 0.20 | 0.32 | 0.26 | 0.13 | 0.19 |
| **39** | 0.16 | 0.13 | 0.24 | 0.10 | 0.14 | 0.19 | 0.13 | 0.10 | 0.11 | 0.26 | 0.11 | 0.09 | 0.06 |

**Table 7 (Contd...):** Nei's unbiased measures of genetic similarity (Nei, 1978)
|  | 14 | 15 | 16 | 17 | 18 | 19 | 20 | 21 | 22 | 23 | 24 | 25 | 26 |
| --- | --- | --- | --- | --- | --- | --- | --- | --- | --- | --- | --- | --- | --- |
| **1** |  |  |  |  |  |  |  |  |  |  |  |  |  |
| **2** |  |  |  |  |  |  |  |  |  |  |  |  |  |
| **3** |  |  |  |  |  |  |  |  |  |  |  |  |  |
| **4** |  |  |  |  |  |  |  |  |  |  |  |  |  |
| **5** |  |  |  |  |  |  |  |  |  |  |  |  |  |
| **6** |  |  |  |  |  |  |  |  |  |  |  |  |  |
| **7** |  |  |  |  |  |  |  |  |  |  |  |  |  |
| **8** |  |  |  |  |  |  |  |  |  |  |  |  |  |
| **9** |  |  |  |  |  |  |  |  |  |  |  |  |  |
| **10** |  |  |  |  |  |  |  |  |  |  |  |  |  |
| **11** |  |  |  |  |  |  |  |  |  |  |  |  |  |
| **12** |  |  |  |  |  |  |  |  |  |  |  |  |  |
| **13** |  |  |  |  |  |  |  |  |  |  |  |  |  |
| **14** | 1.00 |  |  |  |  |  |  |  |  |  |  |  |  |
| **15** | 0.28 | 1.00 |  |  |  |  |  |  |  |  |  |  |  |
| **16** | 0.44 | 0.35 | 1.00 |  |  |  |  |  |  |  |  |  |  |
| **17** | 0.24 | 0.18 | 0.29 | 1.00 |  |  |  |  |  |  |  |  |  |
| **18** | 0.36 | 0.28 | 0.36 | 0.29 | 1.00 |  |  |  |  |  |  |  |  |
| **19** | 0.23 | 0.07 | 0.17 | 0.60 | 0.12 | 1.00 |  |  |  |  |  |  |  |
| **20** | 0.12 | 0.11 | 0.07 | 0.32 | 0.07 | 0.52 | 1.00 |  |  |  |  |  |  |
| **21** | 0.12 | 0.21 | 0.17 | 0.39 | 0.12 | 0.55 | 0.61 | 1.00 |  |  |  |  |  |
| **22** | 0.40 | 0.15 | 0.10 | 0.13 | 0.23 | 0.15 | 0.21 | 0.10 | 1.00 |  |  |  |  |
| **23** | 0.38 | 0.18 | 0.13 | 0.20 | 0.25 | 0.18 | 0.28 | 0.18 | 0.53 | 1.00 |  |  |  |
| **24** | 0.19 | 0.23 | 0.04 | 0.11 | 0.16 | 0.18 | 0.35 | 0.13 | 0.33 | 0.33 | 1.00 |  |  |
| **25** | 0.23 | 0.28 | 0.18 | 0.10 | 0.18 | 0.08 | 0.16 | 0.08 | 0.17 | 0.25 | 0.39 | 1.00 |  |
| **26** | 0.15 | 0.33 | 0.21 | 0.23 | 0.23 | 0.14 | 0.16 | 0.14 | 0.05 | 0.04 | 0.22 | 0.35 | 1.00 |
| **27** | 0.26 | 0.33 | 0.37 | 0.32 | 0.32 | 0.22 | 0.20 | 0.30 | 0.12 | 0.18 | 0.18 | 0.22 | 0.25 |
| **28** | 0.23 | 0.28 | 0.23 | 0.24 | 0.36 | 0.08 | 0.07 | 0.12 | 0.10 | 0.25 | 0.08 | 0.23 | 0.15 |
| **29** | 0.24 | 0.23 | 0.34 | 0.28 | 0.29 | 0.23 | 0.32 | 0.28 | 0.23 | 0.36 | 0.25 | 0.29 | 0.21 |
| **30** | 0.36 | 0.28 | 0.30 | 0.24 | 0.52 | 0.12 | 0.12 | 0.12 | 0.23 | 0.19 | 0.13 | 0.30 | 0.27 |
| **31** | 0.18 | 0.28 | 0.18 | 0.24 | 0.23 | 0.28 | 0.12 | 0.17 | 0.10 | 0.19 | 0.25 | 0.23 | 0.35 |
| **32** | 0.17 | 0.21 | 0.23 | 0.18 | 0.23 | 0.12 | 0.07 | 0.12 | 0.10 | 0.13 | 0.13 | 0.28 | 0.20 |
| **33** | 0.26 | 0.25 | 0.20 | 0.16 | 0.33 | 0.13 | 0.04 | 0.08 | 0.26 | 0.15 | 0.21 | 0.33 | 0.16 |
| **34** | 0.15 | 0.09 | 0.09 | 0.07 | 0.09 | 0.09 | 0.13 | 0.09 | 0.20 | 0.22 | 0.22 | 0.27 | 0.11 |
| **35** | 0.04 | 0.21 | 0.15 | 0.12 | 0.04 | 0.21 | 0.14 | 0.21 | 0.00 | 0.00 | 0.16 | 0.22 | 0.26 |
| **36** | 0.04 | 0.21 | 0.10 | 0.12 | 0.10 | 0.15 | 0.14 | 0.15 | 0.06 | 0.10 | 0.16 | 0.22 | 0.26 |
| **37** | 0.04 | 0.23 | 0.08 | 0.07 | 0.04 | 0.13 | 0.28 | 0.23 | 0.11 | 0.09 | 0.26 | 0.31 | 0.29 |
| **38** | 0.27 | 0.26 | 0.21 | 0.13 | 0.07 | 0.16 | 0.15 | 0.16 | 0.21 | 0.23 | 0.28 | 0.27 | 0.19 |
| **39** | 0.21 | 0.07 | 0.16 | 0.22 | 0.12 | 0.26 | 0.20 | 0.16 | 0.21 | 0.17 | 0.17 | 0.21 | 0.19 |

**Table 7 (Contd...):** Nei's unbiased measures of genetic similarity (Nei, 1978)
|  | 27 | 28 | 29 | 30 | 31 | 32 | 33 | 34 | 35 | 36 | 37 | 38 | 39 |
| --- | --- | --- | --- | --- | --- | --- | --- | --- | --- | --- | --- | --- | --- |
| 1 |  |  |  |  |  |  |  |  |  |  |  |  |  |
| 2 |  |  |  |  |  |  |  |  |  |  |  |  |  |
| 3 |  |  |  |  |  |  |  |  |  |  |  |  |  |
| 4 |  |  |  |  |  |  |  |  |  |  |  |  |  |
| 5 |  |  |  |  |  |  |  |  |  |  |  |  |  |
| 6 |  |  |  |  |  |  |  |  |  |  |  |  |  |
| 7 |  |  |  |  |  |  |  |  |  |  |  |  |  |
| 8 |  |  |  |  |  |  |  |  |  |  |  |  |  |
| 9 |  |  |  |  |  |  |  |  |  |  |  |  |  |
| 10 |  |  |  |  |  |  |  |  |  |  |  |  |  |
| 11 |  |  |  |  |  |  |  |  |  |  |  |  |  |
| 12 |  |  |  |  |  |  |  |  |  |  |  |  |  |
| 13 |  |  |  |  |  |  |  |  |  |  |  |  |  |
| 14 |  |  |  |  |  |  |  |  |  |  |  |  |  |
| 15 |  |  |  |  |  |  |  |  |  |  |  |  |  |
| 16 |  |  |  |  |  |  |  |  |  |  |  |  |  |
| 17 |  |  |  |  |  |  |  |  |  |  |  |  |  |
| 18 |  |  |  |  |  |  |  |  |  |  |  |  |  |
| 19 |  |  |  |  |  |  |  |  |  |  |  |  |  |
| 20 |  |  |  |  |  |  |  |  |  |  |  |  |  |
| 21 |  |  |  |  |  |  |  |  |  |  |  |  |  |
| 22 |  |  |  |  |  |  |  |  |  |  |  |  |  |
| 23 |  |  |  |  |  |  |  |  |  |  |  |  |  |
| 24 |  |  |  |  |  |  |  |  |  |  |  |  |  |
| 25 |  |  |  |  |  |  |  |  |  |  |  |  |  |
| 26 |  |  |  |  |  |  |  |  |  |  |  |  |  |
| 27 | 1.00 |  |  |  |  |  |  |  |  |  |  |  |  |
| 28 | 0.37 | 1.00 |  |  |  |  |  |  |  |  |  |  |  |
| 29 | 0.35 | 0.14 | 1.00 |  |  |  |  |  |  |  |  |  |  |
| 30 | 0.32 | 0.36 | 0.19 | 1.00 |  |  |  |  |  |  |  |  |  |
| 31 | 0.22 | 0.30 | 0.19 | 0.18 | 1.00 |  |  |  |  |  |  |  |  |
| 32 | 0.36 | 0.42 | 0.14 | 0.28 | 0.28 | 1.00 |  |  |  |  |  |  |  |
| 33 | 0.24 | 0.26 | 0.07 | 0.33 | 0.33 | 0.31 | 1.00 |  |  |  |  |  |  |
| 34 | 0.15 | 0.09 | 0.21 | 0.21 | 0.09 | 0.26 | 0.23 | 1.00 |  |  |  |  |  |
| 35 | 0.31 | 0.10 | 0.28 | 0.04 | 0.22 | 0.21 | 0.11 | 0.11 | 1.00 |  |  |  |  |
| 36 | 0.11 | 0.15 | 0.23 | 0.15 | 0.22 | 0.15 | 0.05 | 0.11 | 0.38 | 1.00 |  |  |  |
| 37 | 0.23 | 0.08 | 0.20 | 0.13 | 0.08 | 0.23 | 0.15 | 0.29 | 0.31 | 0.40 | 1.00 |  |  |
| 38 | 0.20 | 0.12 | 0.32 | 0.12 | 0.16 | 0.11 | 0.18 | 0.31 | 0.20 | 0.14 | 0.28 | 1.00 |  |
| 39 | 0.25 | 0.16 | 0.37 | 0.12 | 0.21 | 0.11 | 0.13 | 0.19 | 0.33 | 0.20 | 0.23 | 0.30 | 1.00 |

**Table 8:** Ewens-Watterson test for neutrality for each RAPD’s locus

| Locus | n | k | Allele Frequency* |  |  |  |  |
| --- | --- | --- | --- | --- | --- | --- | --- |
|  |  |  | Obsrv. | Expected |  |  |  |
|  |  |  |  | Mean | SE | L95 | U95 |
| B73-1 | 39 | 2.00 | 0.7413 | 0.7725 | 0.0252 | 0.5003 | 0.9500 |
| B73-2 | 39 | 2.00 | 0.7843 | 0.7749 | 0.0245 | 0.5030 | 0.9500 |
| B73-3 | 39 | 2.00 | 0.7413 | 0.7683 | 0.0258 | 0.5030 | 0.9500 |
| B73-4 | 39 | 2.00 | 0.8294 | 0.7704 | 0.0261 | 0.5003 | 0.9500 |
| B73-5 | 39 | 2.00 | 0.7205 | 0.7749 | 0.0255 | 0.5030 | 0.9500 |
| B73-6 | 39 | 2.00 | 0.7413 | 0.7685 | 0.0260 | 0.5003 | 0.9500 |
| B73-7 | 39 | 2.00 | 0.9494 | 0.7681 | 0.0258 | 0.5003 | 0.9500 |
| B73-8 | 39 | 2.00 | 0.7625 | 0.7658 | 0.0258 | 0.5030 | 0.9500 |
| B74-1 | 39 | 2.00 | 0.8762 | 0.7691 | 0.0269 | 0.5030 | 0.9500 |
| B74-2 | 39 | 2.00 | 0.6618 | 0.7587 | 0.0268 | 0.5003 | 0.9500 |
| B74-3 | 39 | 2.00 | 0.7843 | 0.7675 | 0.0248 | 0.5003 | 0.9500 |
| B74-4 | 39 | 2.00 | 0.7205 | 0.7694 | 0.0266 | 0.5003 | 0.9500 |
| B74-5 | 39 | 2.00 | 0.8294 | 0.7650 | 0.0259 | 0.5003 | 0.9500 |
| B74-6 | 39 | 2.00 | 0.8294 | 0.7748 | 0.0248 | 0.5003 | 0.9500 |
| B75-1 | 39 | 2.00 | 0.8294 | 0.7682 | 0.0253 | 0.5003 | 0.9500 |
| B75-2 | 39 | 2.00 | 0.6436 | 0.7681 | 0.0264 | 0.5003 | 0.9500 |
| B75-3 | 39 | 2.00 | 0.6261 | 0.7746 | 0.0250 | 0.5030 | 0.9500 |
| B75-4 | 39 | 2.00 | 0.7205 | 0.7760 | 0.0242 | 0.5030 | 0.9500 |
| B75-5 | 39 | 2.00 | 0.7625 | 0.7693 | 0.0265 | 0.5003 | 0.9500 |
| B75-6 | 39 | 2.00 | 0.5934 | 0.7679 | 0.0247 | 0.5030 | 0.9500 |
| B75-7 | 39 | 2.00 | 0.5197 | 0.7641 | 0.0262 | 0.5030 | 0.9500 |
| B75-8 | 39 | 2.00 | 0.7205 | 0.7703 | 0.0252 | 0.5030 | 0.9500 |
| B75-9 | 39 | 2.00 | 0.8762 | 0.7656 | 0.0262 | 0.5030 | 0.9500 |
| B76-1 | 39 | 2.00 | 0.9745 | 0.7706 | 0.0254 | 0.5030 | 0.9500 |
| B76-2 | 39 | 2.00 | 0.5120 | 0.7630 | 0.0250 | 0.5003 | 0.9500 |
| B76-3 | 39 | 2.00 | 0.6808 | 0.7715 | 0.0263 | 0.5030 | 0.9500 |
| B76-4 | 39 | 2.00 | 0.5784 | 0.7694 | 0.0257 | 0.5030 | 0.9500 |
| B76-5 | 39 | 2.00 | 0.6618 | 0.7675 | 0.0258 | 0.5030 | 0.9500 |
| B76-6 | 39 | 2.00 | 0.7413 | 0.7723 | 0.0256 | 0.5003 | 0.9500 |
| B76-7 | 39 | 2.00 | 0.6436 | 0.7688 | 0.0258 | 0.5030 | 0.9500 |
| B76-8 | 39 | 2.00 | 0.6436 | 0.7674 | 0.0259 | 0.5003 | 0.9500 |
| B76-9 | 39 | 2.00 | 0.8294 | 0.7642 | 0.0257 | 0.5003 | 0.9500 |
| B76-10 | 39 | 2.00 | 0.9745 | 0.7682 | 0.0259 | 0.5030 | 0.9500 |
| B77-1 | 39 | 2.00 | 0.6436 | 0.7694 | 0.0254 | 0.5030 | 0.9500 |
| B77-2 | 39 | 2.00 | 0.8066 | 0.7753 | 0.0246 | 0.5030 | 0.9500 |
| B77-3 | 39 | 2.00 | 0.9246 | 0.7599 | 0.0260 | 0.5030 | 0.9500 |
| B77-4 | 39 | 2.00 | 0.6808 | 0.7754 | 0.0252 | 0.5030 | 0.9500 |
| B77-5 | 39 | 2.00 | 0.9002 | 0.7719 | 0.0249 | 0.5030 | 0.9500 |
| B77-6 | 39 | 2.00 | 0.9745 | 0.7670 | 0.0259 | 0.5030 | 0.9500 |
| B78-1 | 39 | 2.00 | 0.9494 | 0.7750 | 0.0250 | 0.5030 | 0.9500 |
| B78-2 | 39 | 2.00 | 0.7625 | 0.7670 | 0.0259 | 0.5030 | 0.9500 |
| B78-3 | 39 | 2.00 | 0.7413 | 0.7713 | 0.0265 | 0.5003 | 0.9500 |
| B78-4 | 39 | 2.00 | 0.6808 | 0.7631 | 0.0257 | 0.5030 | 0.9500 |
| B78-5 | 39 | 2.00 | 0.6093 | 0.7758 | 0.0252 | 0.5003 | 0.9500 |
| B78-6 | 39 | 2.00 | 0.7003 | 0.7717 | 0.0247 | 0.5003 | 0.9500 |
| B78-7 | 39 | 2.00 | 0.7413 | 0.7693 | 0.0254 | 0.5030 | 0.9500 |
| B78-8 | 39 | 2.00 | 0.7003 | 0.7735 | 0.0257 | 0.5003 | 0.9500 |
| B78-9 | 39 | 2.00 | 0.7003 | 0.7757 | 0.0269 | 0.5003 | 0.9500 |
| B78-10 | 39 | 2.00 | 0.9246 | 0.7687 | 0.0256 | 0.5030 | 0.9500 |
| B78-11 | 39 | 2.00 | 0.7843 | 0.7685 | 0.0256 | 0.5030 | 0.9500 |
| B78-12 | 39 | 2.00 | 0.8066 | 0.7658 | 0.0252 | 0.5030 | 0.9500 |

**Table 9:** The three classes of degradation capacity of dyes: highly efficient (>50%), medium (50-30%) and lower in efficiency (<30%) degradation of MG and BPB of isolates of *Pleurotus* species

| Classes | Bromophenol Blue (BPB) |  | Malachite greenG (MG) |  |
| --- | --- | --- | --- | --- |
|  | 5<sup>th</sup> day | 10<sup>th</sup> day | 5<sup>th</sup> day | 10<sup>th</sup> day |
| **High** | #6, #11, #15, #18, #22 | #3, #5, #6, #11, #12, #13, #15, #16, #18, #22, #24, #25, #27, #28, #29, #30, #32, #33, #34, #35, #36, #37, #38, #39 | -- | #6 |
| **Moderate** | #5, #16, #24, #25, #29, #33, #39 | #4, #8, #20, #23, #26, #31 | #6 | #3, #5, #9, #11, #14, #15, #16, #18, #20, #22, #25, #26, #27, #29, #30, #33, #34, #35, #37, #38, #39 |
| **low** | #1, #2, #3, #4, #7, #8, #9, #10, #12, #13, #14, #17, #19, #20, #21, #23, #26, #27, #28, #30, #31, #32, #34, #35, #36, #37, #38 | #1, #2, #7, #9, #10, #10, #14, #17, #19, #21 | #1, #2, #3, #4, #5, #7, #8, #9, #10, #11, #12, #13, #14, #15, #16, #17, #18, #19, #20, #21, #22, #23, #24, #25, #26, #27, #28, #29, #30, #31, #32, #33, #34, #35, #36, #37, #38, #39 | #1, #2, #4, #7, #8, #10, #12, #13, #17, #19, #21, #23, #24, #28, #31, #32, #36 |

### Clustering of isolates based on DNA profiles

Dendrogram as illustrated in figure (Fig. 2) was generated by Nei’s genetic distance given in table 7. Each leaf represents an individual observation. The leaves are spaced evenly along the horizontal axis. The vertical axis indicates a distance or dissimilarity measure. The height of a node represents the distance of the two clusters that the node joins. The obtained dendrogram depicts that all isolates fall into two distinct groups (similarity >12%). Similarity indices were developed on the basis of amplified fragments of the 39 different genotypes using 6 RAPD primers (Table 4). The genetic similarity values ranged from 0.36 to 0.93 with a mean of 0.64. All isolates diverged into two major clusters, represented as I and II (Fig. 2) except isolate #06 that clustered separately from all the isolates. The first (cluster-I) major cluster was divisible into two sub-clusters (A & B) at 18% similarity level, in which one sub-cluster (i.e. sub-cluster-A) comprised of three isolates (#35, #36, #37) while the other sub-cluster (sub-cluster-B) again divisible into two sub-sub-cluster (a & b), each comprised four isolates (sub-sub-cluster a-#29, #39, #34, #38 and sub-sub-cluster b-#22, #23, #24, #25).

Second major cluster (cluster-II) was divisible into three sub-clusters (A, B & C). Sub-cluster-A comprised eight isolates (#31, #26, #32, #28, #27, #33, #30 & #18), sub-cluster-B comprised ten isolates (#15, #11, #16, #14, #9, #13, #5, #12, #10 & #3), while sub-cluster-C further divisible into two sub-sub-clusters-a & b. Sub-sub-clusters-a comprised four isolates (#21, #20, #19 & #17), while sub-sub-clusters-b comprised five isolates (#8, #7, #4, #2 and #1).

**Figure 2:**
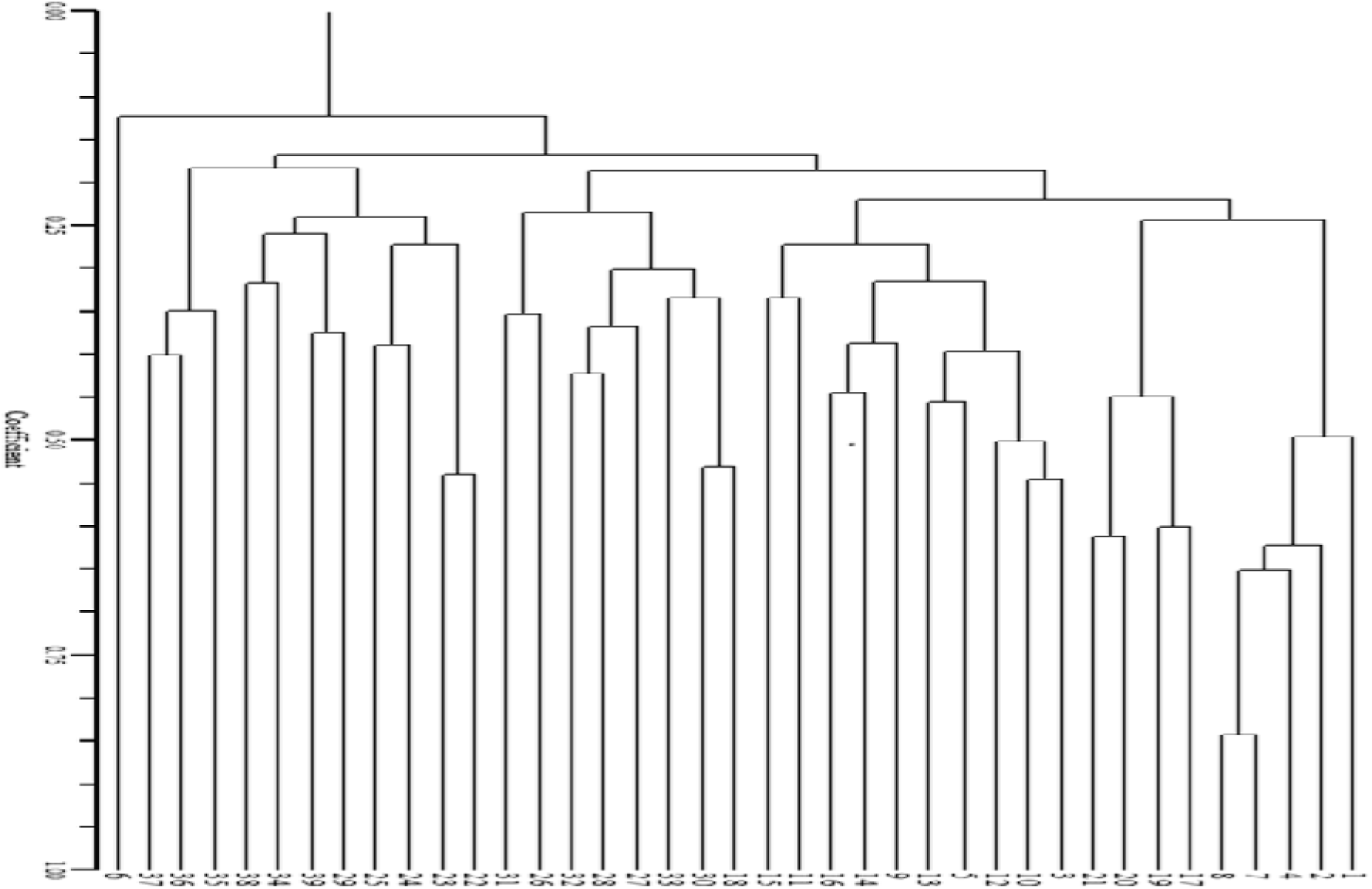
Dendrogram prepared from genetic similarity matrices obtained through RAPD analysis by using UPGMA of 39 isolates of *Pleurotus*

### Availability of nutraceuticals

Total crude protein quantified in dried fruiting bodies of different isolate of *Pleurotus* species is given in figure 3. The highest amount of protein was observed in #29 i.e., 28.48 mg/100mg when compared with 39 isolates; and the lowest content is in #31 with a value of 17.16 mg/100mg. The amount of protein present in any strain is dependent on many factors basically types of substrate and other additional ingredients of substrate including weather conditions such as temperature. The total carbohydrate value was calculated and found in the range from 3.55 to 5.43g/100g of fresh oyster mushroom as given in figure 4. From figure (Fig. 4), it was observed that the highest carbohydrate content (5.43 g/100g) was found in #10, however, lowest in #18. The total phenolic content was determined in the fresh fruiting body of isolates of *Pleurotus* collected in the present study and draw a bar diagram as depicted in figure 5. The content of phenolics were different in different isolates and it was in range of 21.19 to 36.32 mg/g, in which #21 showed maximum and #01 showed minimum phenolic content of in their fruiting body.

**Figure 3:**
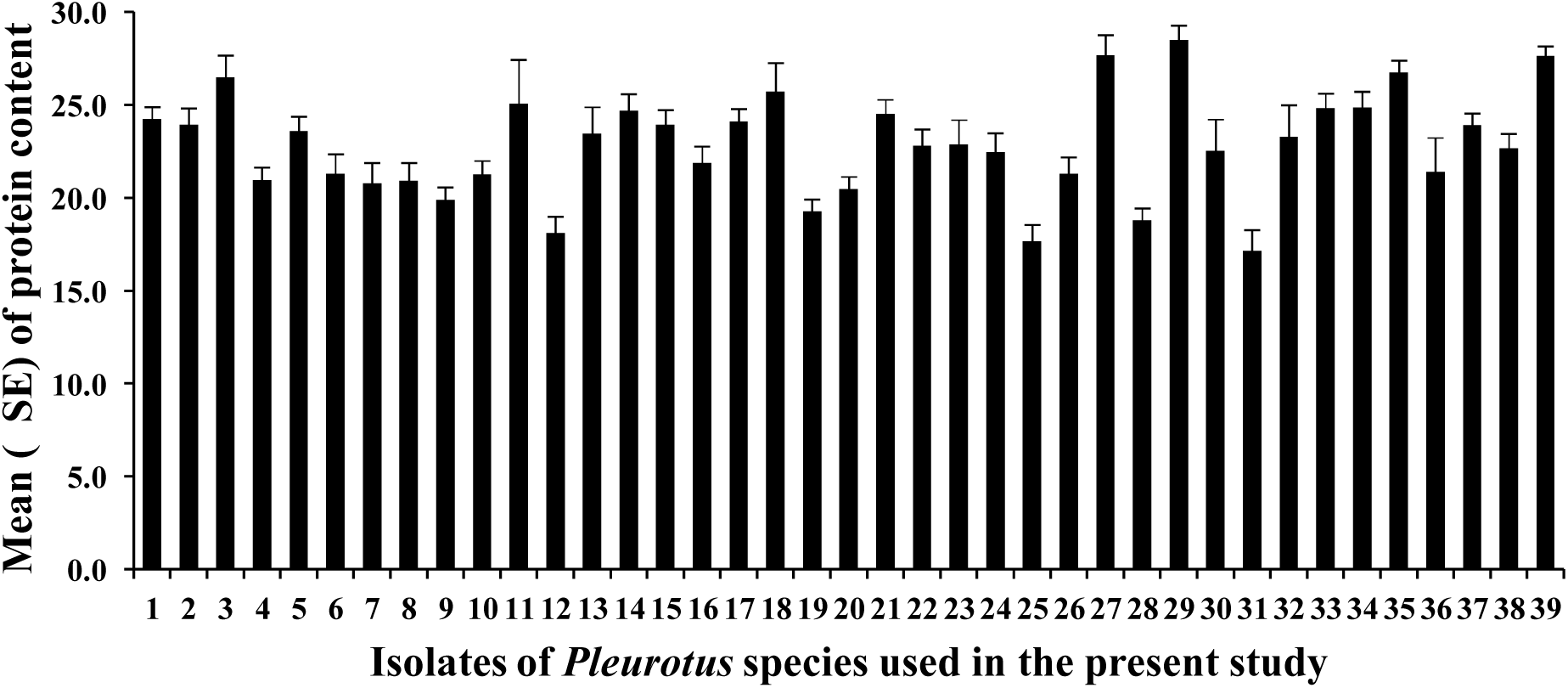
Content of total proteins measured by Lowry et al. (1951) method

**Figure 4:**
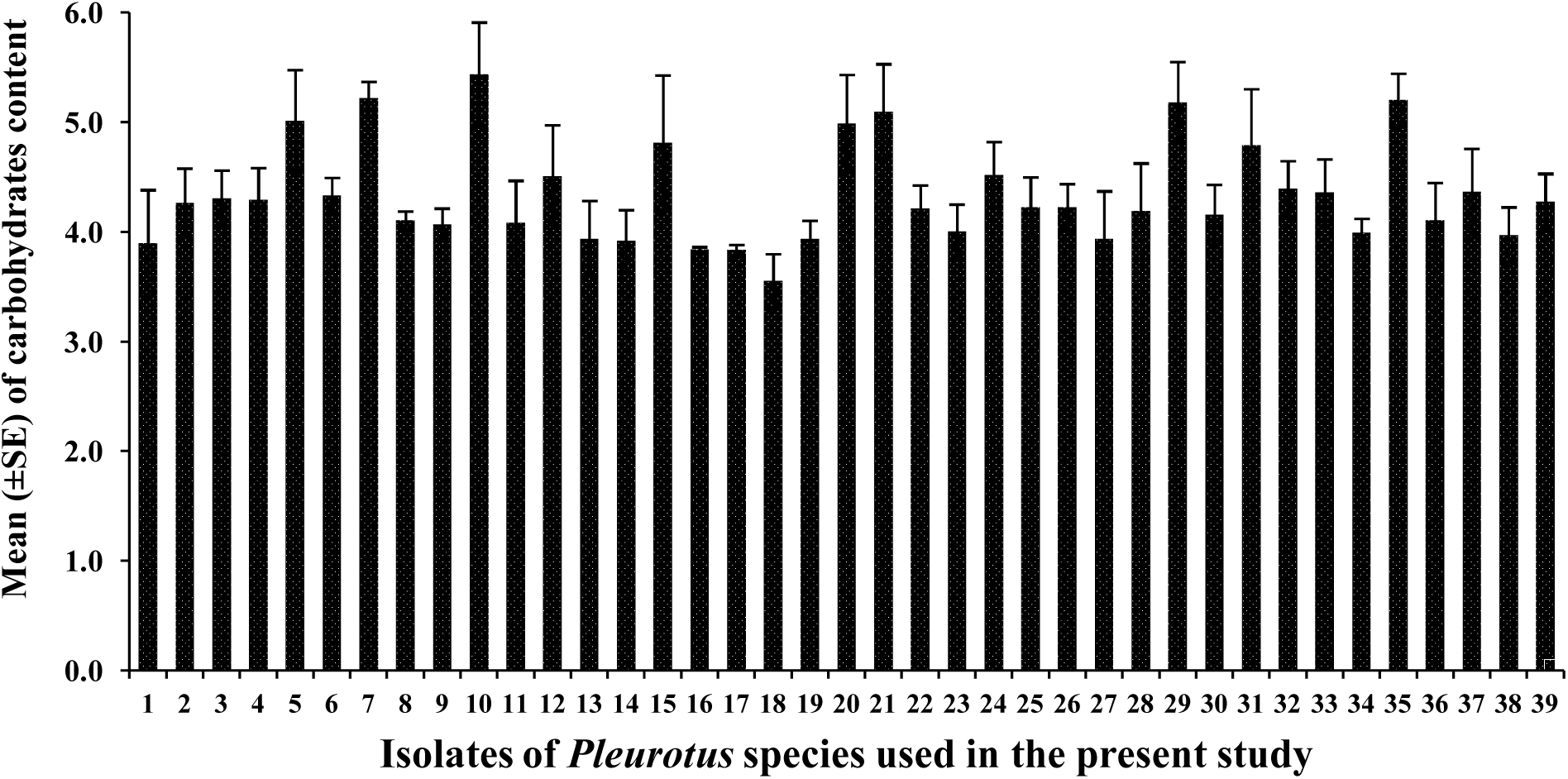
Content of total carbohydrates measured by phenol sulphuric acid method

**Figure 5:**
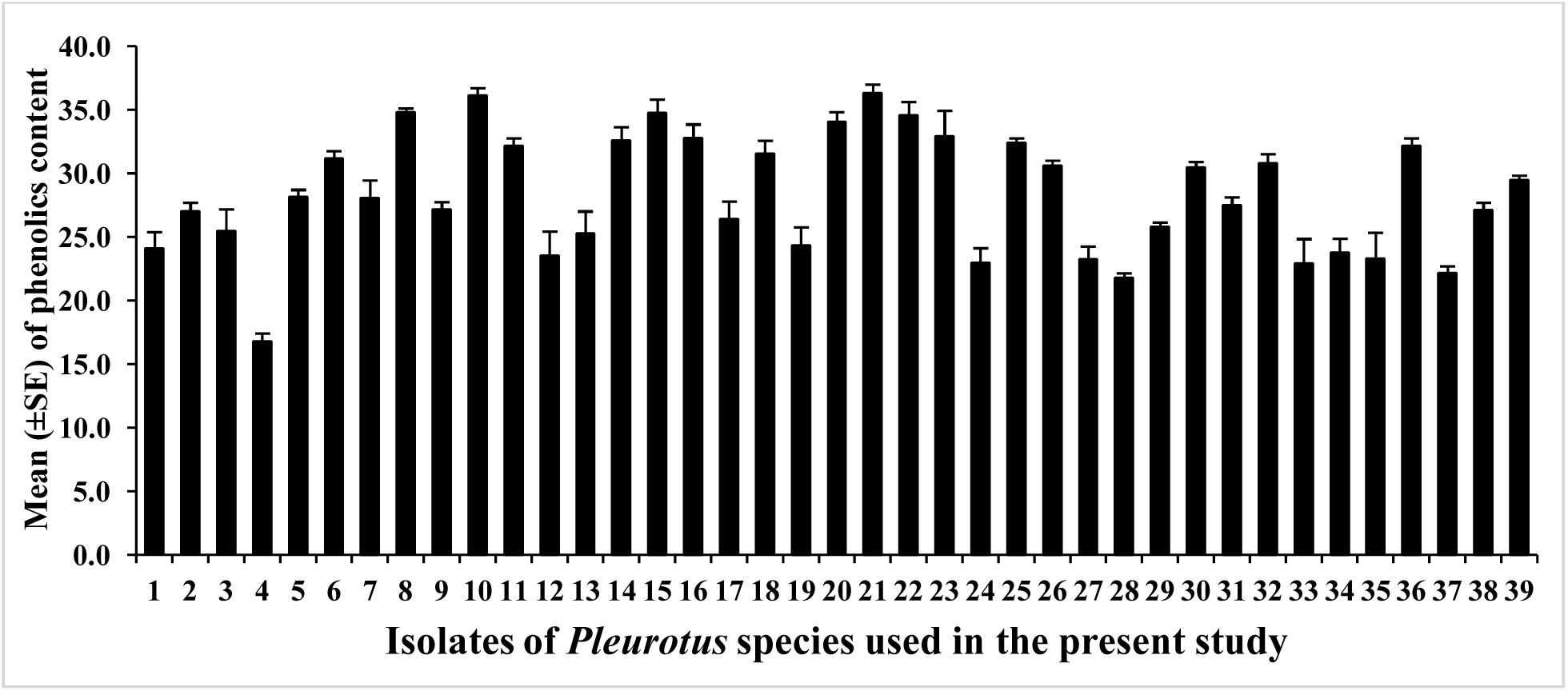
Content of total phenolic measured by Singleton et al. (1965) method

Vitamin B_12_ content of different isolates was determined by bioassay with vitamin B_12_ requiring microorganisms, such as *Lactobacillus delbrueckii* subsp. *lactis* as described by Schneider (1987). The level of vitamin B_12_ was assayed in different wild isolates of *Pleurotus* spp. A good amount of vitamin B_12_ was observed, which was in range of 0.05 to 0.32mg/kg (of dried mushroom) of vitamin B_12_ in different isolates of *Pleurotus* spp as given in figure (Fig. 6). From **figure 6**, highest amount of vitamin B_12_ is observed in isolate #32 i.e., 0.32 mg/kg and lowest in #36.

**Figure 6:**
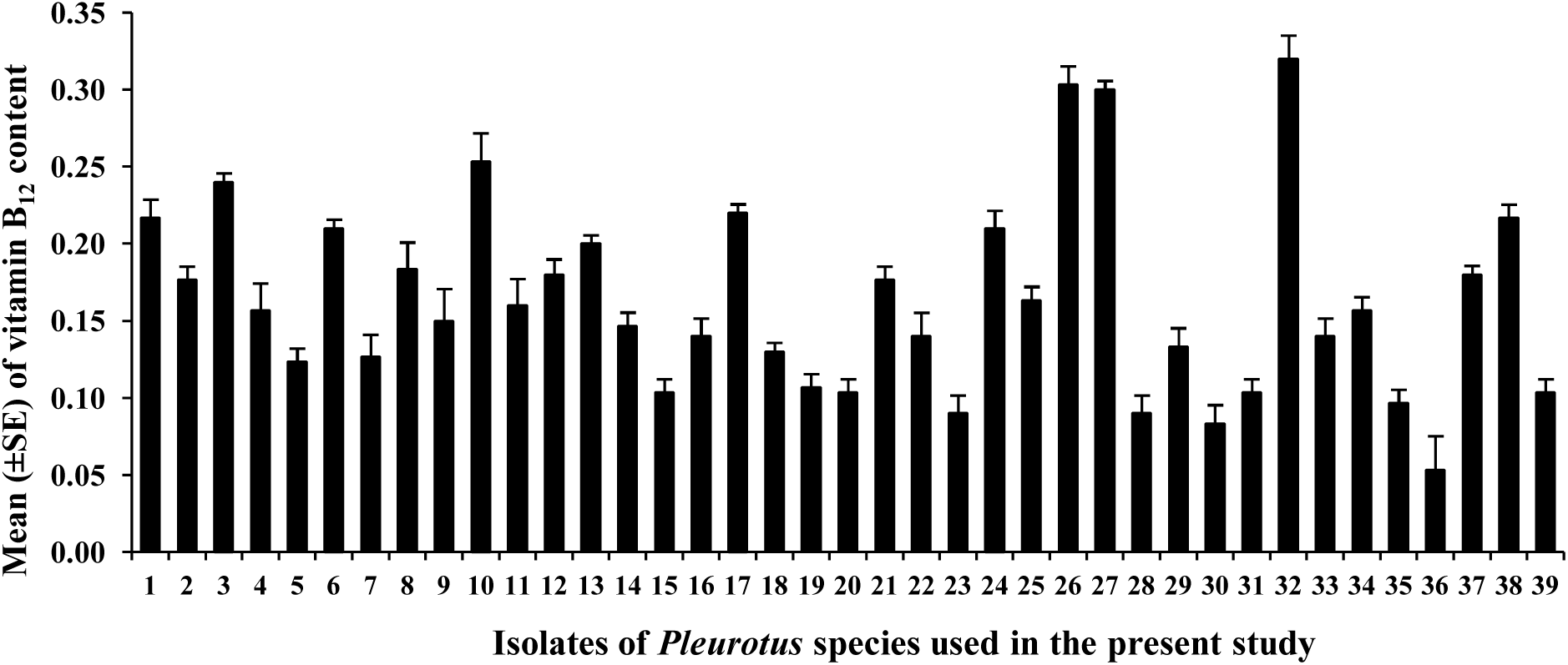
Vitamin B_12_ contents in fruiting bodies of different isolates of *Pleurotus* species

### Laccase enzyme

Production of fungal laccase was assayed via the oxidation of Guaiacol (o-methoxyphenol catechol monomethylether) as substrate as per the method used by Arora and Sandhu (1985) by the extracellular enzyme obtained through liquid state fermentation from growing mycelia of different isolates of *Pleurotus* spp. Figure 7 showed that #2 produced maximum and #27 minimum laccase enzyme i.e., 4.03 to 19.13 IU/ml respectively.

**Figure 7:**
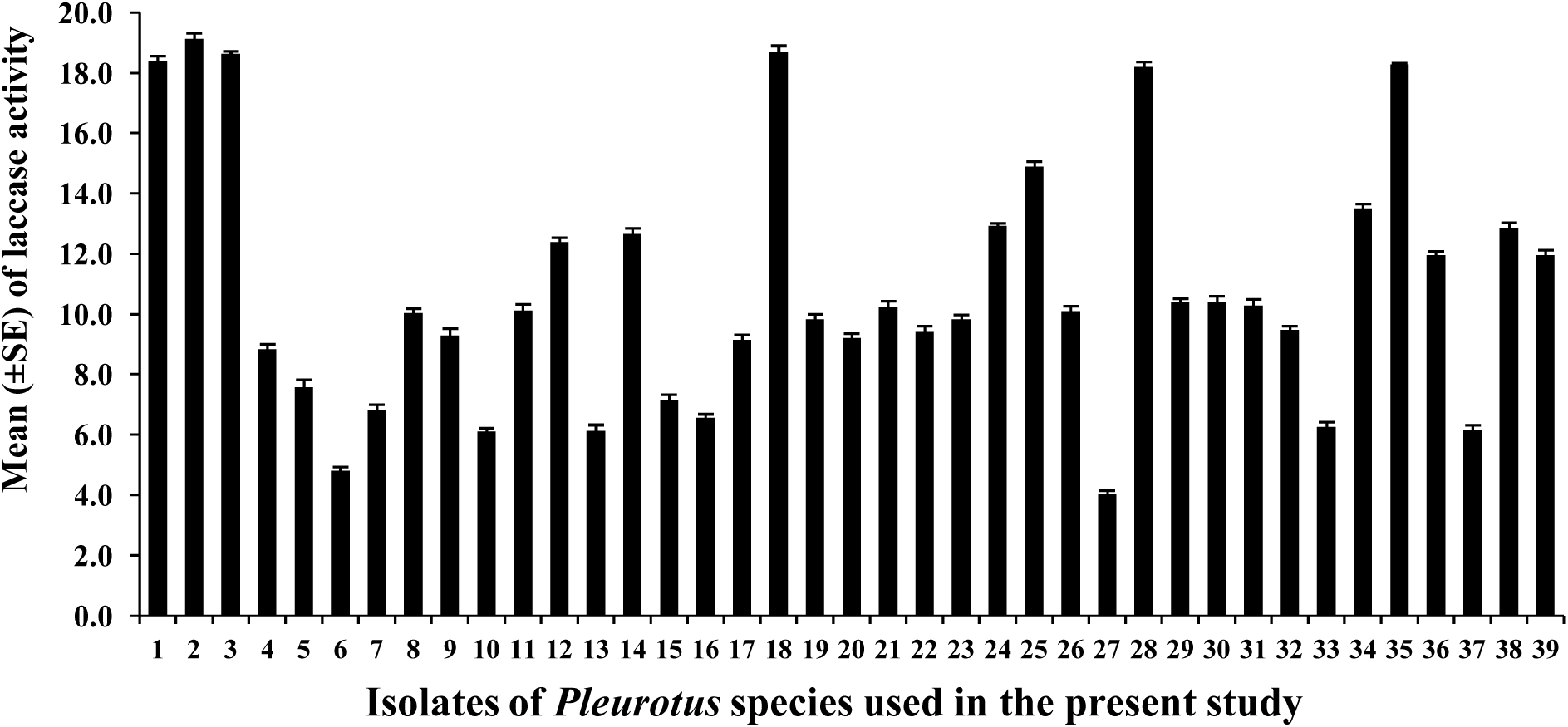
Laccase enzyme activity assayed by oxidation of Guaiacol (o-methoxyphenol catechol monomethylether) as substrate as per the method used by Arora and Sandhu (1985)

### Decolorization of malachite greenG (0.01%) and bromophenol blue (0.05%)

The *Pleurotus* isolates were analyzed for their decolorizing ability of two textile dyes i.e., malachite greenG (MG) and bromophenol blue (BPB) on solid medium. The isolates were able to decolorize MG (in terms of decolorized area) within 5-10 days of incubation at 28±1°C when PDA was supplemented with 0.01% (w/v) of MG. from the observation of figure 8, most of the isolates were not efficient to complete decolorization of MG. Only isolate #06 showed highest ability of degradation of MG dye (>75%). however rest isolates showed lower capacity in removal of MG. Isolate #06 decolorized >50% within 5 days. Similarly, decolorization potential for BPB was observed as given in figures (Fig. 9 and 10). The decolorization of dye was observed by the bleaching activity observed under and around of developing mycelia.

**Figure 8:**
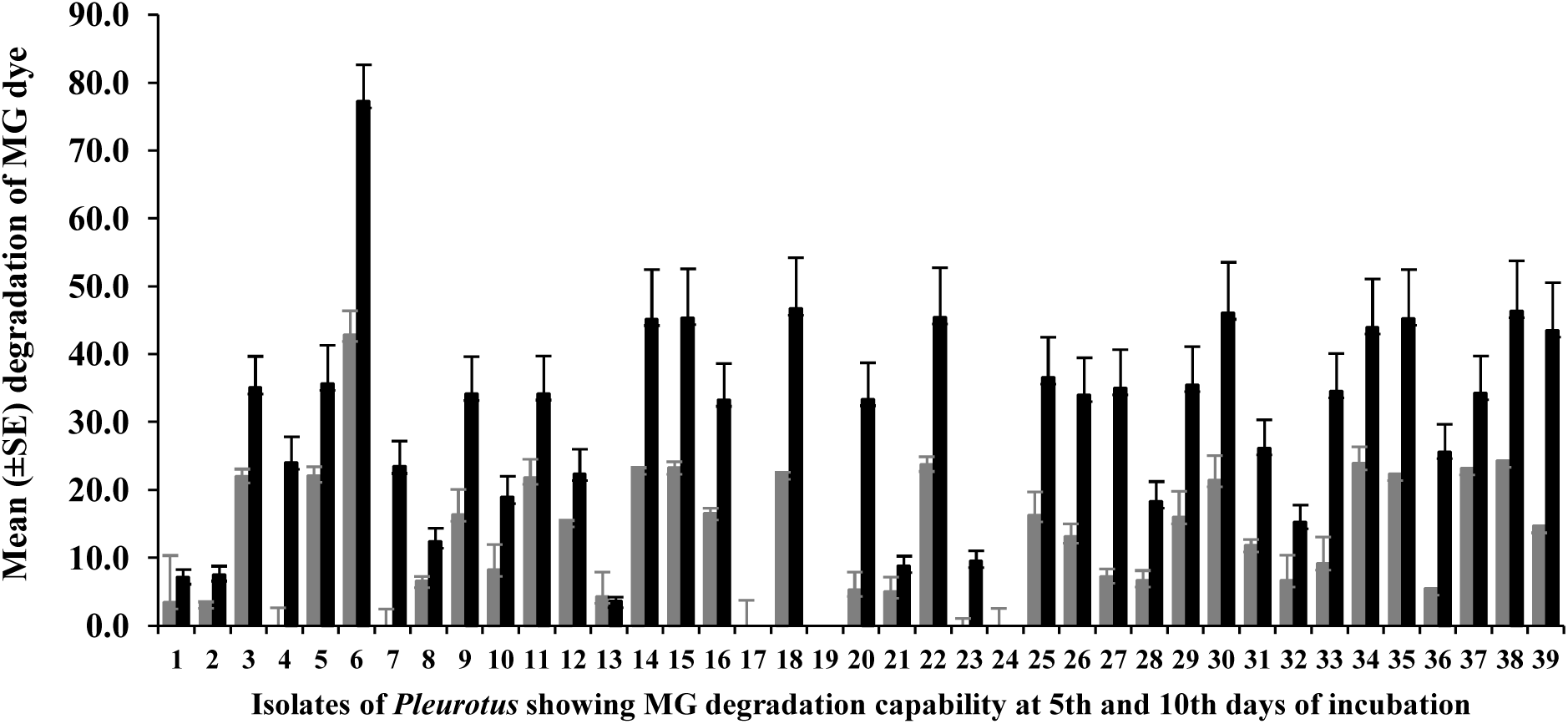
Isolates of *Pleurotus* species showing decolorization of malachite greenG (0.01% (w/v) concentration at 5^th^ day and 10^th^ day of incubation at 28±°C

**Figure 9:**
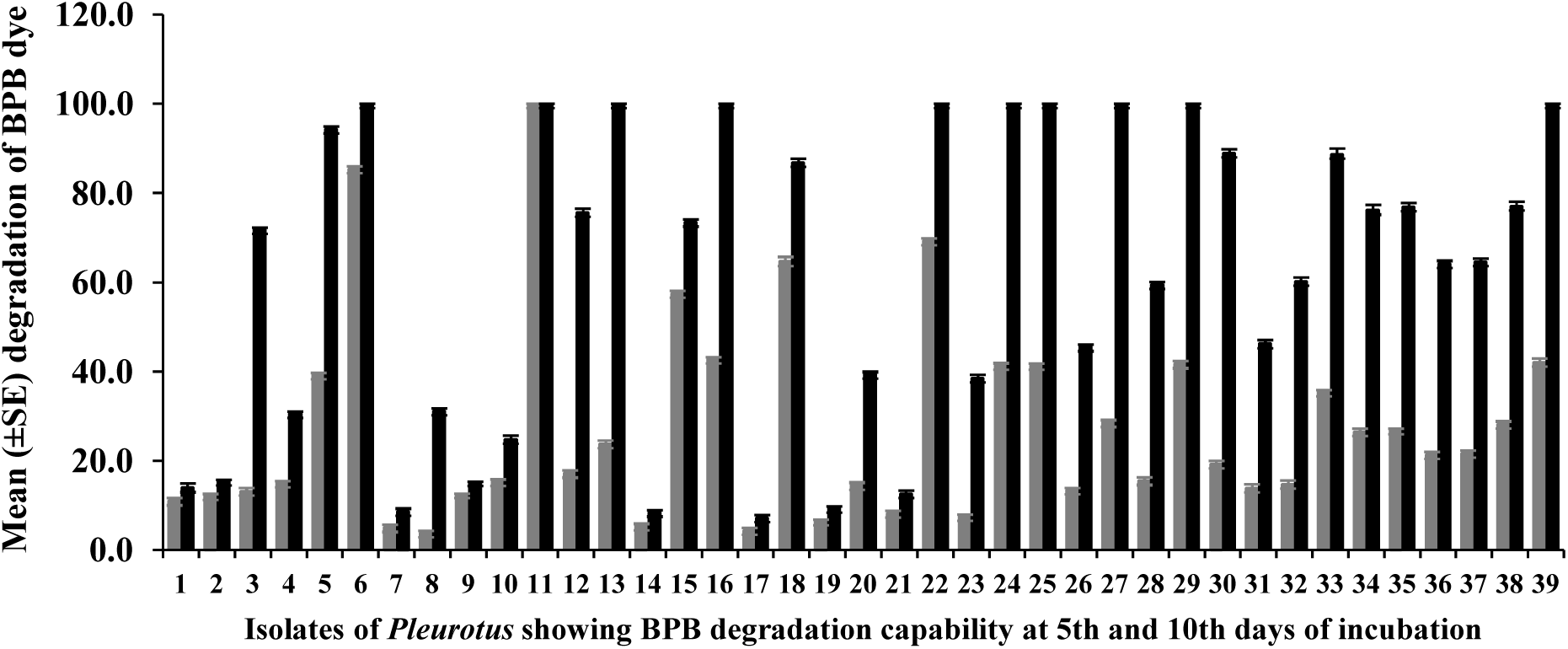
Isolates of *Pleurotus* species showing decolorization of Bromophenol blue (0.05% (w/v) concentration at 5^th^ day and 10^th^ day of incubation at 28±°C

## Discussion

In the present study, RAPD markers showed more effective discrimination of closely related species of *Pleurotus* (Fig. 2), out of 10 arbitrary primers used for the RAPD profiling of the all isolates, 6 primers exhibited a total of 51 polymorphic bands as given in table (Table 4), and all were 100% polymorphic bands. Number of amplified products (bands) varied depending upon the primers used; primers B-73, B-74, B-75, B-76, B-77 and B-78 generated 8, 6, 9, 10, 6, and 12 bands, respectively. These variations in the number of bands may be due to the sequence of primer, availability of annealing sites in the genome and template quality (Kernodle et al., 1993). Afzal et al. (2004) reported only 75% polymorphism while studying 21 cultivars of mungbean employing 34 decamer primer while in another study 73.2% polymorphism was reported in an indigenous medicinal herb *Gymnema sylvestre* with 15 decamer primers (Nair and Keshavachandran, 2006), the later obtained an average of 6.0 bands per primer. The result 8.5 bands per primer of this study is similar to finding of Afzal et al. (2004), however, Yin et al. (2012) reported a still higher average value (about 13.7). Chandra et al. (2010) used the RAPD markers and found variations in banding pattern to discriminate the eight *Pleurotus* species and concluded that every isolate of this study was characterized by a different RAPD genotype. Unlike MLEE, RAPD analyses generally detect occurrence of a single allele that assesses polymorphisms at a wide range of loci (William et al., 1990). Later, many researchers (Dowdy and McGaughey, 1996; Pornkulwat et al., 1998) differentiated very closely related species or even geographical populations using RAPDs. Use of RAPDs provided a much clearer picture of the relationships between *Pleurotus* species and studied eco-geographic zones. Different number of bands was generated depending upon the primers used, and for the sake of convenience the result of RAPD profile of all the 39 isolates was categorized into three group *viz.* high, moderate and no polymorphism. As per the results given earlier in table (Table 4), some primers (B-78, B-76, B-75 and B-73) generated high polymorphisms while B-77 and B-74 primers exhibited moderate polymorphisms. By comparing their banding pattern it was found that fifty one putative loci were resolved with sufficient consistency and clarity. Gene frequency was found in the range of 0.012 to 0.987, in which the highest gene frequency was 0.422 of allele-1 for *B76-2* loci while the lowest gene frequency was 0.012 of allele-1 for *B77-6* loci. Alleles having frequency less than 0.01 were not detected. In certain pair of alleles (e.g. *B73-2, B74-3* & *B78-2*; *B76—3* & *B77-4*) frequencies of alleles are uniform that may indicate the active participation of natural selection in maintaining genetic polymorphisms as discussed.

In order to understand the genetic diversity more critically, more biometric parameters like Effective number of alleles, Nei’s (1973) genetic distance and Shannon’s information index were also calculated. Summary of results of Nei’s (1973) gene diversity and Shannon information index is given in table 6. Genetic diversity was found from 0.025 to 0.488 whereas, the mean 0.244±0.116; which indicates that many of the loci differ between all pairs of RAPD genotypes. Lowest gene diversity was shown by *B76-1* loci with a value of 0.025 while highest was by *B76-2* loci showing a value of 0.488. These results suggested that the RAPD approach showed considerable potential for *Pleurotus* species discrimination. These results are similar to Yin et al. (2012) who reported substantial genetic diversity (0.22 to 0.97) amongst 15 different cultivars of *P. pulmonarious* using RAPD. Similar results (genetic diversity: 0.178-0.262) were obtained by Zervakis et al. (2001), while studying genetic polymorphism in *P. eryngii* species complex growing in greater Mediterranean area. In the present study the observed number of alleles was 2.000±0.00 while effective number of alleles was 1.355±0.21 as given in table (Table 6); and the Shannon’s Information Index varied from 0.1205 to 0.6811 with an average 0.397±0.155. These results are in agreement with Boldo et al. (2003) who made epidemiological studies with 47 clinical and reference strains of *Candida glabrata* from several geographical origins. The value of this index is related to diversity of isolates, value <0.5 is considered to be of low diversity while value >0.5 indicates higher diversity. Studies from Howell et al. (1994) also emphasized that RAPD technique is far more reliable for identification analysis than morphological and physiological identification because in the later methods characteristics are influenced by cultivation conditions as discussed earlier. Coefficients of genetic similarity were calculated from paired comparison of the all 39 isolates, based on normalized identity of each locus in each of species (Nei, 1978). The results are given in table (Table 6), and it was in range from 0.11 to 0.84 similar to the results of Chandra et al. (2010) who used 8 Indian *Pleurotus* species (procured from ICAR-Dirctorate of Mushroom Research, Solan, H.P., India) with 10 decamer random primers to discriminate the genetic divergence of different species.

So far the statistical analyses of data obtained from RAPD profiling are in the form of loci and alleles. Therefore, any polymorphism between two samples based upon RAPD marker is the manifestation of polymorphism of its loci and alleles. It might be expected that RAPD markers would relate with eco-edaphic zones of studied area, from where isolates collected. However, it is to be noted that RAPDs produce dominant markers, and in addition, it scans different parts of the DNA generating large number of markers, and hence, different results may be obtained, such as a RAPD marker reveals the nucleotide differences in a random sequence of DNA of 10 bases long (if a decamer is used). The estimation of genetic variation by this method, as performed in this study with RAPD, permits better estimations of genetic diversity from any species as compared to morphological and physiological parameter.

In order to see the relationship amongst the isolates used in this study, a dendrogram was generated from the pair wise distance matrices. The clustering pattern in the Jaccard’s similarity, dendrogram generated by RAPD as given in figure (Fig. 2) demonstrated discriminating power of RAPDs with reference to their eco-geographic zones. Similar results were obtained by Liu and Furnier (1993) in a study of the genetic variation in aspen (*Populus* spp.), and by Lanner-Herrera et al. (1996) who studied the diversity in natural populations of wild kale, *Brassica oleracea* L. Dendrogram (Fig. 2) generated through RAPD data from UPGMA cluster analysis demonstrated that all isolates clustered into two distinct groups in the distance 0.12. All isolates fell into two major clusters except isolate #06 that come into viewed separately from all the clustered isolates; and all these were grouped into seven clusters.

The statistical analysis based upon the allelic frequencies of RAPDs loci separated all the isolates into seven clusters. At the distance of 0.12, most isolates formed clusters and they are further grouped as discussed earlier. The first cluster mainly comprised five isolates (#1, #2, #4, #7, #8,), second cluster comprised four isolates (#17, #19, #20, #21), third cluster comprised ten isolates (#3, #10, #12, #5, #13, #9, #14, #16, #12, #15), fourth cluster comprised eight isolates (#18, #30, #33, #27, #28, #32, #26, #31), fifth cluster comprised eight isolates (#22, #23, #24, #25, #29, #39, #34, #38), sixth cluster comprised only three isolates (#35, #36, #37) while single isolate (#6) represented seventh cluster. However, it is worth noting that all isolates used in this study almost grouped with reference to their respective eco-geographic zones. Discrimination of isolates by RAPD markers with reference to their respective eco-geographic zones suggested that geographic isolation strongly influenced the evolution of the populations as similarly explained by Sun et al. (1999).

These results indicated some correlation amongst the isolates with respect to their collection site/native place, similar results were observed by Sonnante et al. (1997) who studied the genetic diversity within and between *Vigna luteola* and *V. Marina* (fodder crop). They observed that RAPD markers were able to disclose a much higher level of polymorphisms based upon isozymes profile essentially at the intraspecific level. A possible explanation for the differences found among these dendrograms might be based on the kind of information provided by each type of marker. These RAPDs detect variation in both coding and non-coding regions. Small, repeated, and random sequence mutations would be accumulated in non-coding sequences, and the diversity can be revealed by RAPD. Another factor which needs to be considered for RAPD analysis is that bands of identical mobility may occasionally correspond to non-homologous fragments (Chalmers et al., 1992; Tinker et al., 1993). Although, in an epidemiological study involving pathogenic isolates of *Aspergillus fumigatus,* dendrogram was prepared using isozyme and RAPDs were very coherent on the basis of cophenetic analysis (Rinyu et al., 1995). This could be due to the fact that high value of cophenetic correlation coefficient was due to the high number of negative matches, since on use of the Jaccard’s coefficient, which does not take into account the negative matches, however, in another study involving the population of *Elymus caninus* (a species of flowering plant in the Poaceae family) dendrograms derived from isozyme and RAPD data showed no correlation between clusters and geographic origins (Sun et al., 1999).

It is well accepted that the level of genetic variation is generally considered adaptive and related to the breadth of geographical ranges and/or to the ecological heterogeneity within the ranges (Lewinsohn *et al.,* 2000; Nevo, 1988). Speciation and the development of species richness appear to be facilitated by restricted gene flow and isolation of small populations (Lande, 1984). Hence, the high diversity in many intraspecific taxa that are tropically highly specialized suggests that ecologically specialized populations are particularly prone to speciation (Futuyma, 1986a). However, if those populations are brought into contact, much of the divergence they have accomplished will be lost by interbreeding. On the other hand, if they have evolved in to a new species they can retain their diverse adaptations, and refine them even while sympatric (Futuyma, 1986b). In many cases sympatric populations are in an intermediate stage of speciation (i.e. partially reproductively isolated), and they usually interbreed along a hybrid zone that can persist for long periods (Futuyma, 1986a).

The present study was completed in the eastern part of Uttar Pradesh; commonly called Purvanchal comprises of more than fifteen districts including Allahabad, Azamgarh, Jaunpur, Mirzapur, S.R.N. Bhadohi and Varanasi (plus half a dozen more carved out from above districts). The topology of this region is considerably heterogeneous, with a gradient of temperature, precipitation, water logging (Puri, 1992) which is more suitable for generating diversity. Long- and short-term environmental factors (e.g. flood, drought and soil erosion) and likely are crucial to the creation and maintenance of high biodiversity (Taylor and Skinner, 1998). The diversity of this region is considered endangered due to fragmentation of critical habitat (DellaSala et al., 1999). In this study some sort of association was observed with their geographic origin of isolates with RAPD profiling, however less or no association was observed when MLEE was employed as reported by Patel et al. (2018). Environmental and edaphic factors are known to influence the diversity of terrestrial forms in general (Boddy et al., 2013) that also influence genotype of oyster mushroom fungus in a long run.

The degrees of decolorization of different dyes such as malachite green, indigo carmine, xylidine ponceau, Bismarck brown and methyl orange using the white rot fungus *P. ostreatus* were previously evaluated by various researchers (Cerniglia and Sutherland, 2010). These studies demonstrated the potentialities of white rot fungi in bioremediation of dye contaminated ecosystems. The laccase (Revankar and Lele, 2007) and MnP (Tsukihara et al., 2008) play a major role in complete oxidation of Direct Blue 14 (DB14) (Vishwakarma et al., 2012). It was observed that laccase and peroxidases can act as starters of a chain reaction which leads to dye degradation by generating highly active free radicals (e.g., Mn^3+^, lipid, hydroxyl, and peroxy-radicals) (Hofrichter, 2002; Rabinovich et al., 2004). According to Meyer (Meyer, 1981), because of the structural variety of azo dyes, they are not uniformly susceptible to biodegradation. It was demonstrated that substituent groups such as nitro and sulpho are frequently recalcitrant to biodegradation, whereas 2 methyl, 2-methoxy, 2, 6-dimethyl and 2,6-dimethoxy-substituted 4-(4-sulfophenylazo)-phenol were preferred for azo-dye degradation by peroxidase from *Streptomyces* spp. and *Phanerochaete chrysosporium* (Suzuki et al., 2001). The breaking down of the dye into smaller fragments, including the breakage of the azo bond, can lead to a decrease in the absorbance of the visible spectra and leave a colorless solution (Zille et al., 2003). White rot fungi degrade lignin because they secrete oxidoreductases including lignin peroxidase (1,2-bis (3,4-dimethoxyphenyl) propene-1,3-diol:hydrogen-peroxide-Lip EC 1.1.1.14), manganese peroxidase (Mn(II): hydrogenperoxide oxidoreductase EC 1.11.1.13), and laccase (benzenediol: oxygen reductase EC 1.10.3.2). These enzymes oxidize in a nonspecific way both phenolic and nonphenolic lignin derivatives and thus are promising candidates for the degradation of environmental pollutants, for example, phenols, anilines, dyes, lignocelluloses (Fahr et al., 1999; Ferreira-Leitão et al., 2007; Vishwakarma et al., 2012] and highly recalcitrant compounds such as polychlorinated biphenyls (PCBs) and polycyclic aromatic hydrocarbons (PAHs) and lignin derivatives has been attributed to the oxidative enzymes, especially laccase (Riccardi et al., 2005).

The results were found similar to the report analyzed by Watanabe et al. (2012) in six edible mushrooms. Surprisingly, we also assayed laccase concentration in liquid mycelia growth medium and it was in accordance of many previous studies (Xie et al., 2016; Inacio et al., 2015). The laccase production could be improved by supplementing in liquid growth medium (Park et al., 2014).

Dried fruiting bodies of different isolates contain sufficient amount of crude protein (Fig. 3) and reported by many researchers also (Tolera and Abera, 2017). Similarly, total carbohydrates and phenolic content measured also, and found very lower carbohydrates content which make it very suitable food for diabetic patient especially (Tolera and Abera, 2017; Widyastuti et al., 2015; Parul and Asha, 2014). Total phenolics in this study were in range 21.19 to 36.32mg/g and it was found in agreement with previous report (Tan et al., 2015; Abugri and McElhenney, 2013). From the figure 4 it is clear that *Pleurotus* have sufficient quantity of vitamin B_12_ to fulfill the daily requirement of our population. Though adults need only 2.3 to 5.0µg of B_12_ per day for optimum health, dietary intake of B_12_ should exceed that amount due to the complex process required to assimilate and metabolize this essential nutrient. Uptake of this vitamin in the gastrointestinal tract depends on intrinsic factor, which is synthesized by the gastric parietal cells, and on the cubam receptor in the distal ileum (Nielsen et al., 2012). As per guidelines of the Institute of Medicine (USA), consumption of approximately 100 g of dried oyster mushrooms could provide the recommended daily dietary allowance (2.4µg/day) for adults (Sullivan and Herbert, 1965).

## Conclusion

Present study demonstrates that the molecular markers generated through RAPD are more useful as compared to morphological markers for evaluating genetic diversity through characterization and identification of relationships among *Pleurotus* species of mushrooms vis-a-vis geographical zones. It indicated a high level of genetic polymorphism amongst the isolates of *Pleurotus* species despite availability of relatively lower number of isolates of *Pleurotus*. The dendrogram based upon RAPDs reflected better geographic affinities that took into account all DNA fragments. Although no evidence of selective effects of any polymorphic loci was recorded in this study because the correlation with climatic and physical variables were non-significant. Hence, the isolates of different zones are meaningfully addressed by the dendrograms obtained from RAPD data, which correlated and discriminated eco-geographic group by RAPD markers suggests that geographic isolation may influence the evolution of the populations. The Oyster mushroom studied in terms of diversity were also evaluated for their many potentials including protein, carbohydrates, vitamin B_12_, laccase enzyme, degradation of textile dyes and phenolic; which showed variability amongst isolates. The results of this study indicate the potential of diversity in terms of natural products.

## Acknowledgments

Author thanks to Veer Bahadur Singh Purvanchal University, Jaunpur (UP), India, for financial and lab support to this work.

